# Exosomes and exosomal miRNAs from muscle-derived fibroblasts promote skeletal muscle fibrosis

**DOI:** 10.1101/267963

**Authors:** Simona Zanotti, Sara Gibertini, Flavia Blasevich, Cinzia Bragato, Alessandra Ruggieri, Simona Saredi, Clelia Introna, Pia Bernasconi, Lorenzo Maggi, Renato Mantegazza, Marina Mora

## Abstract

We investigated in vitro and in vivo the pro-fibrotic role of exosomes released by muscle-derived fibroblasts of Duchenne muscular dystrophy (DMD) patients, and of miRNAs carried by exosomes. We found that exosomes from DMD fibroblasts, but not from myoblasts, had significantly higher levels of miR-199a-5p, a miRNA up-regulated in fibrotic conditions, compared to control exosomes. In control fibroblasts, exposure to DMD fibroblast-derived exosomes induced a myofibroblastic phenotype with increase in α-smooth actin, collagen and fibronectin transcript and protein expression, soluble collagen production and deposition, cell proliferation, and activation of Akt and ERK signalling, while exposure to control exosomes did not. These findings were related to transfer of high levels of miR-199a-5p and to reduction of its target caveolin-1. Finally, injection of DMD fibroblast-derived exosomes into mouse tibialis anterior muscle after cardiotoxin-induced necrosis, produced greater fibrosis than control exosomes.

Our findings indicate that exosomes produced by local fibroblasts in the DMD muscle are able to induce phenotypic conversion of normal fibroblasts to myofibroblasts thereby increasing the fibrotic response; and suggest miR-199a-5p and caveolin-1as potential therapeutic targets.

## Introduction

Repair normally occurs following acute muscle injury as result of elimination of necrotic fibers by inflammatory cells and replacement of wasted tissue by activation of the myogenic program (Best & Hunter, 2000). However, in chronic severe conditions, such as in Duchenne muscular dystrophy (DMD) and congenital muscular dystrophies, muscle tissue damage and inflammatory infiltrates persist, resident satellite cells are unable to keep pace with the requirement for new fibres, and fibrogenic progenitor cells are continuously activated, leading to massive connective tissue deposition (fibrosis) (Tidball, 2011). Fibrosis can therefore be considered a degenerated tissue repair process. In addition to DMD and chronic myopathies, fibrosis characterizes diseases of various tissues, including lung (Selman *et al*, 2001), liver, kidney, heart, skin, and bladder (Wynn, 2007). Cellular and molecular mechanisms, primarily involving myofibroblasts and transforming growth factor-β1 (TGF-β1), are common to all these conditions. Myofibroblasts, in particular, are key mediators of fibrosis: when activated by a variety of conditions they function as the primary source of collagen accumulation (Kramann *et al*, 2013). In chronic injury, these cells exhibit greater proliferation and increased resistance to apoptosis, both aspects resulting in dysregulation of extracellular matrix (ECM) homeostasis. TGF-β1, on the other hand, is the most potent driver of tissue fibrosis: it induces synthesis of several ECM components and stimulates sustained fibroblast-to-myofibroblast differentiation (Piersma *et al*, 2015; Thannickal *et al*, 2003).

Different types of cells, depending on the injured tissue, may differentiate into myofibroblasts. These include mesenchymal, epithelial and endothelial cells, such as circulating cells from bone marrow, haematopoietic precursors, macrophage subpopulations, smooth muscle cells, pericytes and local fibroblast pools (Kalluri & Weinberg, 2009; Kramann *et al*, 2013). In the fibrotic process fibroblasts play a key role by secreting proteins of the ECM and growth factors, and by differentiating into contractile and secretory myofibroblasts, characterized by α-smooth muscle actin (α-SMA) expression and active ECM synthesis (Hinz *et al*, 2012). Previous in vitro studies by our group have shown that primary fibroblasts derived from DMD muscles have indeed a pro-fibrotic phenotype (Zanotti *et al*, 2010 and 2011) and that several ECM components are altered in DMD myotubes (Zanotti *et al*, 2007). These findings indicate that the fibrotic process of DMD skeletal muscle involves both resident fibroblasts and myogenic cells.

MicroRNAs (miRNAs) are small non-coding RNA molecules found in plants and animals, whose main function is to down-regulate target genes. MiRNAs play important roles in several biological and pathological processes. Recently they have emerged as regulators of cell-cell communications and as paracrine signal mediators (Bang *et al*, 2014; Fujita *et al*, 2014; Kohlhapp *et al*, 2015). Circulating miRNAs are actively transported by RNA-binding proteins or by exosomes (Valadi *et al*, 2007; Zomer *et al*, 2010).

Aberrant expression of miRNAs has been implicated in the development of fibrosis in various tissues through regulation of anti- and pro-fibrotic genes (Bowen *et al*, 2013). Recent studies have shown aberrant expression of several miRNAs, called “fibromiRs”, during fibrosis. These include miR-27a-3p, that negatively regulates lung fibrosis by targeting myofibroblast differentiation (Cui *et al*, 2016), miR-125b critical for fibroblast to myofibroblast transition in cardiac fibrosis (Nagpal *et al*, 2016), and miR-132 in renal fibrosis (Bijkerk *et al*, 2016). MiR-9-5p, another miRNA implicated in fibrosis has been demonstrated to significantly delay TGF-β1-dependent transformation of dermal fibroblasts when over-expressed, and to decrease collagen secretion and the expression of ECM proteins, and α-SMA (Miguel *et al*, 2016). We have recently demonstrated that miR-21 and miR-29 are dysregulated in DMD and that these miRNAs play opposing roles in regulating fibrosis in skeletal muscle (Zanotti *et al*, 2015).

Exosomes are endosomal-derived membrane vesicles (40-100 nm in diameter) secreted in the extracellular space by most cell types. They are natural carrier systems, transporting mRNAs, miRNAs, proteins and lipids between donor and recipient cells, thus actively contributing to cell-cell communication (Mathivanan *et al*, 2010). Exosomes are delimited by a lipid bilayer and contain a common conserved set of proteins and several others that vary depending on cell type, defining a unique tissue/cell type exosome signature, as also reported in freely available databases such as ExoCarta (http://www.exocarta.org) and Vesiclepedia (http://microvesicles.org/index.html). Most common proteins found in exosomes are tetraspannins, such as CD63, specific stress proteins (heat shock proteins: HSPs) such as HSP70, and components of the endosomal sorting complexes required for transport (ESCRT), such as Alix.

In the present study we investigated exosome biogenesis and the mechanisms of exosome release by fibroblasts derived from muscle biopsies of DMD patients, and of exosome uptake by fibroblasts derived from muscle biopsies of control subjects, and evaluated the effects of the horizontal transfer of exosomal content from DMD to control fibroblasts. By focused microarray analysis, we demonstrate that exosomes isolated from DMD fibroblasts have a pro-fibrotic miRNA profile expressing, in particular, high level of miR-199a-5p. We also show that fibroblast-derived DMD exosomes are able to induce a fibrotic phenotype in control fibroblasts and that miR-199a-5p, conveyed by exosomes, is sufficient to induce transdifferentiation of control fibroblasts to myofibroblasts. Finally, we show significantly greater extent of fibrosis in the tibialis anterior muscle of mice injected with DMD fibroblast-derived exosomes (hereinafter called DMD exosomes) than in the contralateral muscle injected with control fibroblast-derived exosomes (hereinafter called control exosomes).

## Results

### Characterization of exosomes

Immunostaining with prolyl-4-hydroxylase to evaluate purity of the muscle-derived fibroblast population obtained after immunomagnetic selection, showed that all cells were positively stained both in control and DMD fibroblasts (Figure 1A).

**Figure 1.**
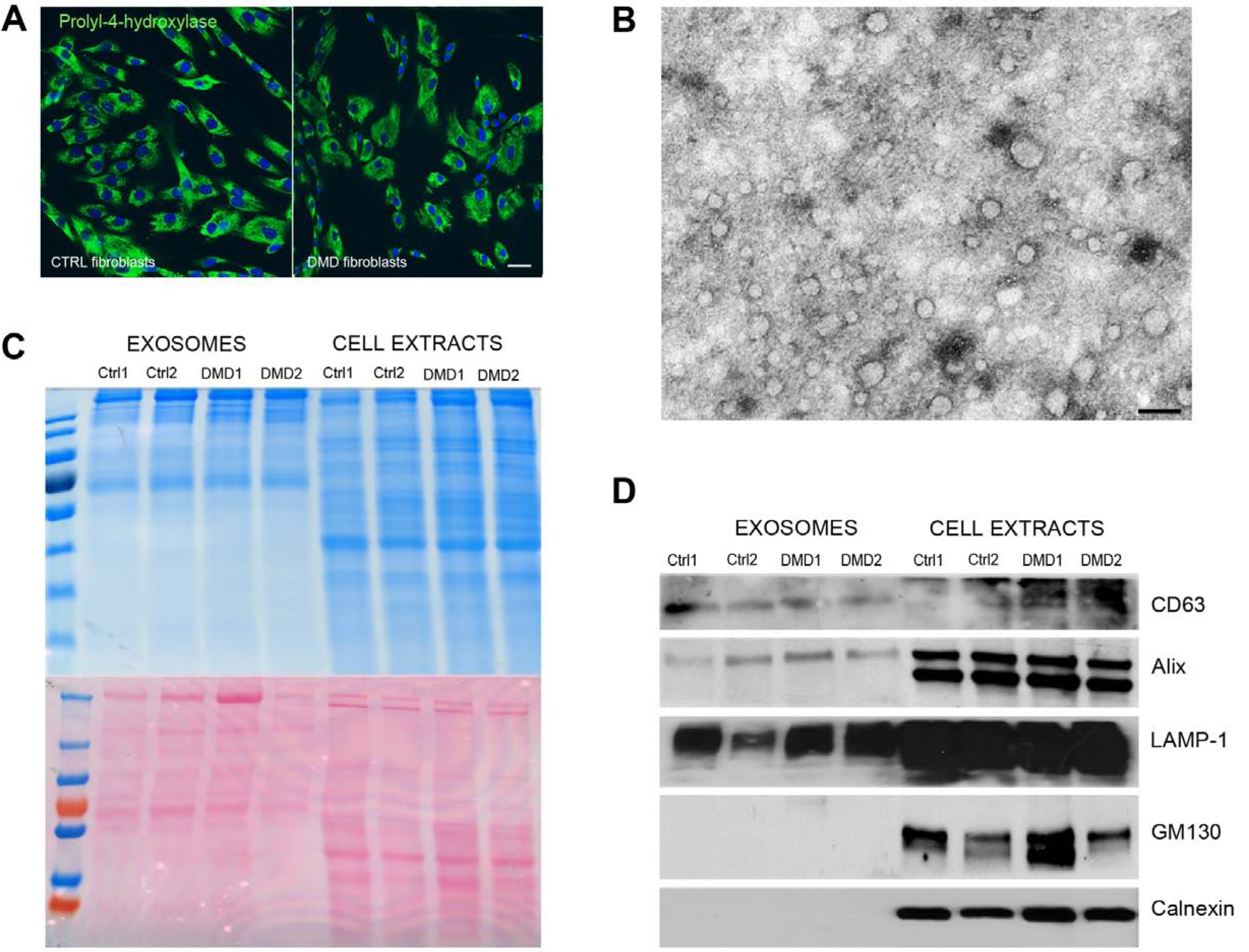
Exosome characterization. **A** Prolyl-4-hydroxylase staining confirming fibroblast purity after immunomagnetic selection in both control and DMD cell cultures. Cell nuclei were stained with DAPI. Scale bar: 10 μm. **B** Electron microscopy showing exosomes as small round vesicles of approximately 30–80 nm in diameter. Scale bar: 100 nm. **C** Page Blue (upper) and Ponceau (lower) staining of exosomal proteins and total cell extracts from two control fibroblast cell lines and two DMD fibroblasts cell lines. **D** Western blot of control and DMD exosomal proteins and cell extracts showing positivity of exosomes for CD63, Alix and LAMP-1, but not for GM130, and calnexin, markers of contaminating cellular proteins.

Staining of exosomal and cell extract protein content with PageBlue and Ponceau S, respectively on gels and nitrocellulose membranes, showed a markedly different protein profile in the lanes loaded with exosomes compared to the lanes loaded with extracts of donor cells (Figure 1C). Notably, the protein profile was identical for control and DMD exosomes derived from fibroblasts or myoblasts.

Western blot analysis confirmed presence of the exosomal markers CD63 and Alix and absence of the cellular markers calnexin (endoplasmic reticulum) and GM130 (Golgi), in the exosomal fraction, indicating that no contaminating cellular debris was present in exosome preparations (Figure 1D). When observed under the electron microscope, exosomes isolated from cell culture media of control and DMD fibroblasts appeared as small round vesicles of approximately 30−80 nm in diameter consistent with reported exosome morphology and size (Figure 1B).

### Cellular exosomal up-take and localization

In order to evaluate whether exosomes could be taken up by recipient cells, the isolated exosomes were labelled with PKH67 dye and then added to the culture media of control fibroblasts. When cells were exposed to the labelled exosomes a green fluorescence punctate signal was observed after 2 h in the cytoplasm and around the nuclei, becoming more evident after 4 h, and markedly evident after 24 h (Figure 2A). No signal was observed in cells treated with unlabelled exosomes. Exosome internalization into the cells was confirmed by colocalization of PKH67-labelled exosomes with α-tubulin, a microtubule marker (Figure 2B) and by differential interference contrast (DIC) microscopy (Figure 2C). To exclude that exosome internalization was a passive transfer, paraformaldehyde-fixed fibroblasts were incubated with PKH67 labelled-exosomes, and no exosomes were observed in the cytoplasm after 4 h exposure at 37°C (not shown). Cells exposed to labelled exosomes and incubated at 4°C showed a very weak signal (Figure 2D), indicating that uptake was an energy-dependent process. The MTT assay, performed after exposure of fibroblasts for 48 h to different exosome concentrations, indicated that no major cytotoxicity effects were induced in the cells (Figure 2E).

**Figure 2.**
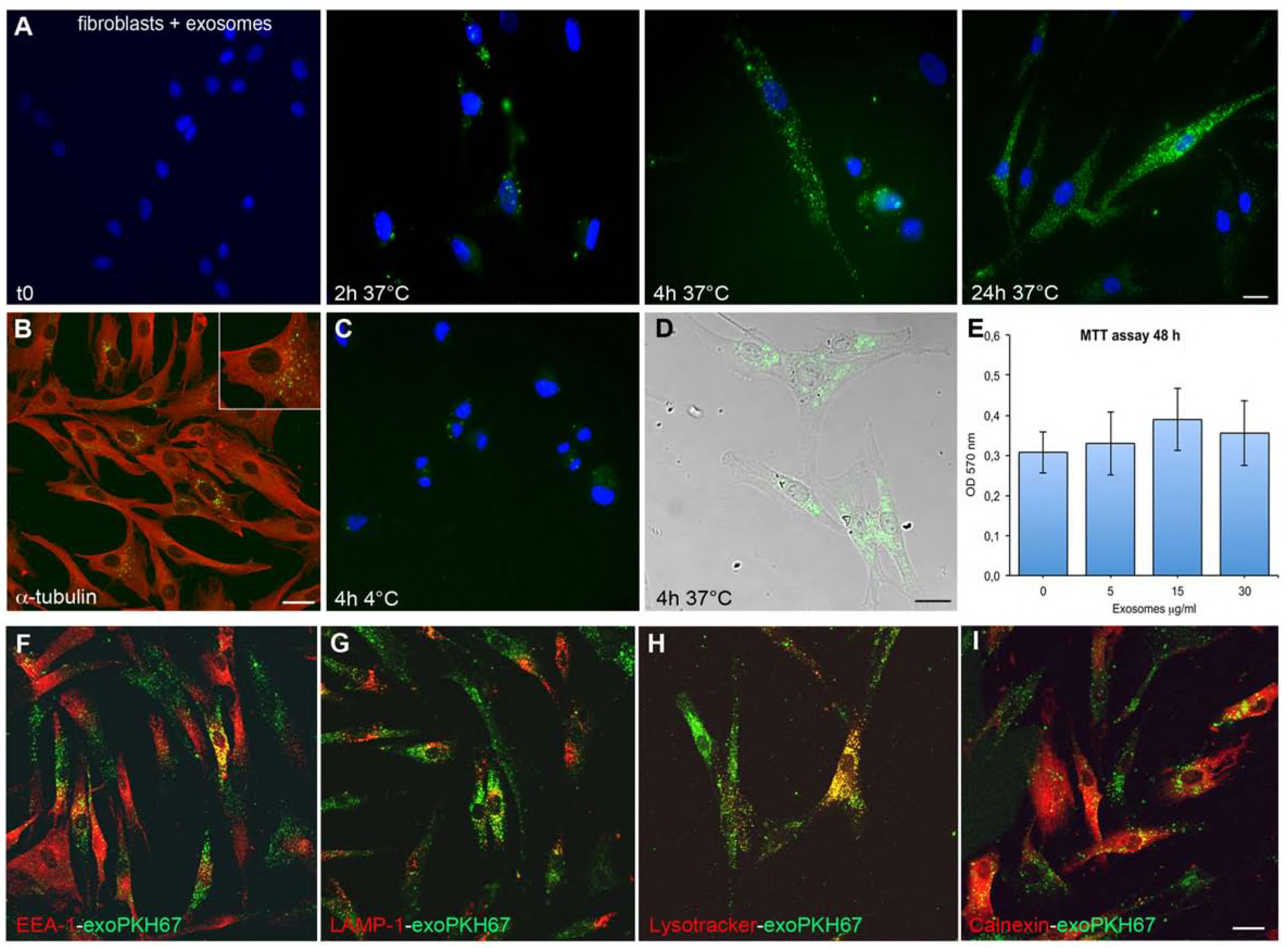
Time course of exosomal uptake and localization. **A** Time course of PKH67-labelled exosome (green) uptake by control fibroblasts showing increasing punctate fluorescence in the cytosol over time. **B** Co-localization of control PKH67-labelled exosomes and α-tubulin (red) showing exosome internalization after 4 h of incubation; a detail is magnified in the inset. Scale bar: 10 μm. **C** Image of control fibroblasts incubated for 4 h at 4°C with PKH67-labelled exosomes showing very limited exosome uptake. Cell nuclei were stained with DAPI. Scale bar: 10 μm. **D** Nomarski interference contrast image of control fibroblasts showing green punctate staining in the cytosol especially near nuclei after 4 h of incubation. Scale bar: 10 μm. **E** MTT assay performed in control fibroblasts after 48 h of exposure to different concentrations of control exosomes showing lack of cytotoxicity. **F-I** Images of control fibroblasts with PKH67-labelled exosomes (green) co-localized with EEA-1 (red), marker of endosomes (F), LAMP-1 (red) (G) and lysotracker (red) (H), both markers of lysosomes, and calnexin (red), marker of endoplasmic reticulum (I), showing partial colocalization with the three first markers but not with the fourth. Scale bar: 10 μm.

Immunocytochemistry showed partial co-localization of exosomes with endosomes (Figure 2F) and lysosomes (Figure 2G,H), but not with the endoplasmic reticulum (Figure 2I) suggesting that, after uptake and internalization into the cells, exosomes diffused within the cytoplasm targeting the endocytic pathway.

### Exosome uptake requires cytoskeletal integrity and is not mediated by fibronectin, and exosome release is ceramide-dependent

Cell treatment with cytochalasin B, a drug that strongly inhibits cytoskeletal network formation by disrupting actin filaments, blocked exosome uptake. In particular, in cytochalasin B-treated cells the uptake of labelled exosomes was significantly reduced (1.699 ± 1.255 vs. 12.01 ± 5.84; p=0.139×10^−5^) compared to untreated cells (for both conditions a total of 380 cells was counted). Addition of the vital dye CellTracker Blue CMAC, (after 4 h of cytochalasin B treatment) to test cell viability excluded that the reduced uptake was due to cell death (Figure 3A,B).

**Figure 3.**
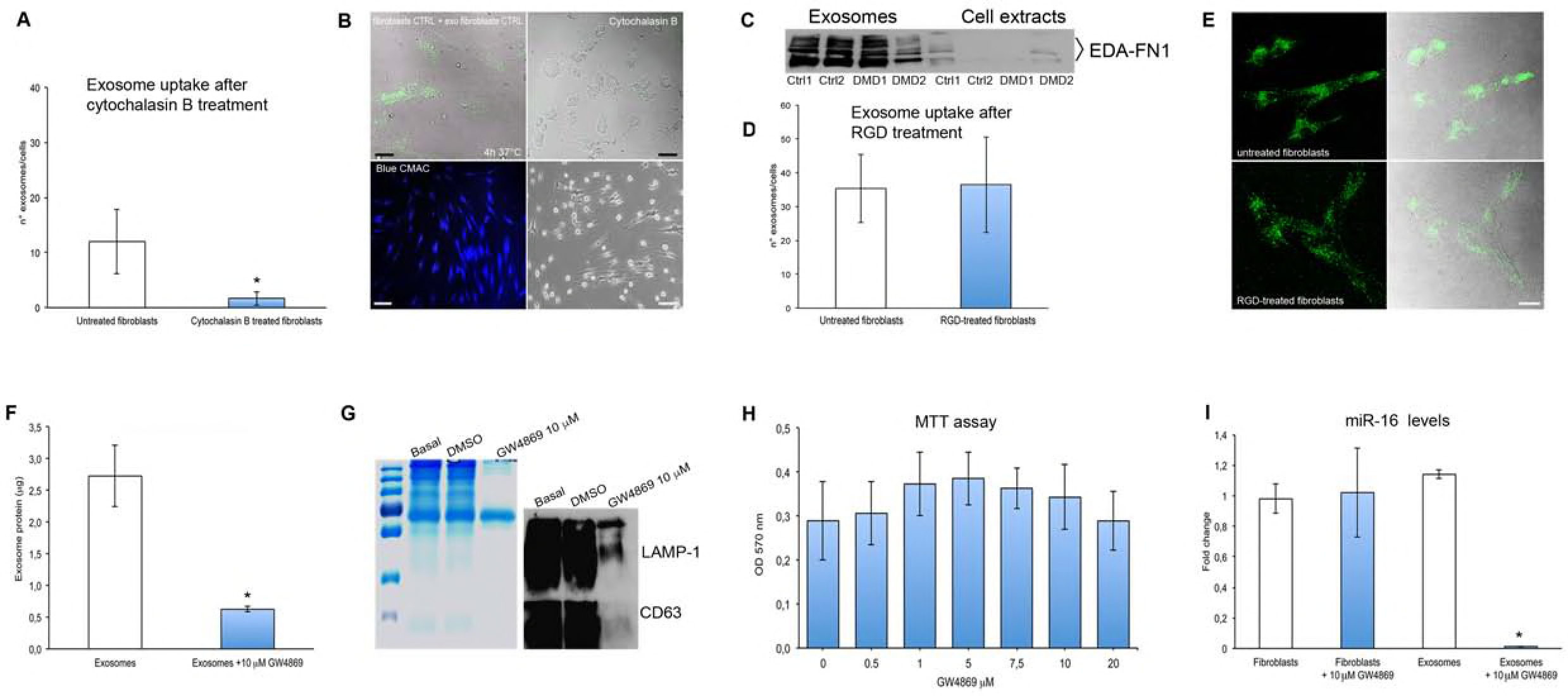
Exosome uptake and release. **A** Cellular uptake of exosomes after cytochalasin B treatment compared to untreated fibroblasts. **B** Nomarski interference contrast of control fibroblasts exposed to PHK67-lebelled exosomes (green) before and after cytochalasin B treatment (upper panels). Scale bar: 20 μm. Bright field of vital dye Cell tracker Blue CMAC staining (lower left panel) showing that during cytochalasin B treatment, despite the morphological alterations due to actin disruption (lower right panel) cells are vital. Scale bar: 50 μm. **C** Representative western blot showing presence of intensely positive EDA-FN1 bands, both in control and DMD fibroblast-derived exosomes, and a weak signal in cell extracts. **D** Exosome uptake after treatment with RGD peptide showing no significant difference between treated and untreated fibroblasts. **E** Confocal and Nomarski interference contrast (NIC) of RGD treated (lower panels) and untreated fibroblasts (upper panels) incubated with PKH67-labelled exosomes showing no significant difference in exosome uptake. Scale bar: 10 μm. **F** Exosomal protein content after treatment with 10 μM GW4869 for 48 h compared to untreated exosomes. **G** Page Blue staining of exosomes isolated from control fibroblasts, showing significant reduction of exosomal proteins after GW4869 treatment as compared to basal conditions, or DMSO (as vehicle); representative western blot of the same samples showing exosomal markers LAMP-1 and CD63 reduced band intensity in GW4869 treated exosomes. **H** MTT assay showing lack of significant cytotoxicity in fibroblasts treated with different concentrations of GW4869 for 48 h. **I** RT-PCR for miR-16, a housekeeping miRNA, showing significantly reduced miR-16 levels in exosomes isolated from fibroblasts treated with GW4869 compared to exosomes from untreated fibroblasts, and no significant difference between GW4869 treated and untreated fibroblasts. Data are presented as mean ± SD, *n* = 3 per group, and analyzed by 2-tailed Student’s *t* test. P ≤ 0.05.

Western blot analysis of control and DMD exosomes, showed a marked expression of the alternative spliced domain of FN1 (EDA-FN1) (Figure 3C). EDA-FN1 is involved in ECM deposition, cell migration and cell adhesion (To & Midwood, 2011). To investigate whether an integrin-EDA-FN1 interaction could mediate exosome uptake, control and DMD fibroblasts were pre-treated with the RGD peptide, a cell adhesion motif present within fibronectin, which blocks integrin-FN1 binding (Atay *et al*, 2011). PKH67-labelled exosomes were then applied to these cells and counted. Exosome numbers in the cytoplasm of RGD-treated and untreated cells were not significantly different (36.51 ± 14.15 vs. 35.36 ± 10.17, p= 0.678406) (Figure 3 D,E), suggesting that exosome uptake by fibroblasts did not involve EDA-FN1-integrin interaction.

Ceramide has been demonstrated to facilitate the formation of endosomal vesicles and the export of miRNAs to exosomes. To evaluate the role of ceramide in exosome biogenesis and release, muscle-derived fibroblasts were treated with neutral sphingomyelinase 2 (nSMase2-GW4869), an inhibitor of ceramide biosynthesis (Trajcovic *et al*, 2008; Kosaka *et al*, 2013). A significant reduction in exosome production, evaluated as protein content, was observed after GW4869-treatment (0.757 ± 0.153 μg vs. 2.28 ± 0.527 μg; p=0.0001) compared to basal condition (Fig 3F); and confirmed by PageBlue Protein staining (Figure 3G). Furthermore, a reduction in signal intensity of the bands corresponding to the exosomal markers LAMP-1 and CD63 was observed by Western blot in exosomal fractions purified from the supernatant of GW4869-treated fibroblasts, compared with the supernatant of control untreated cells or cells treated with DMSO (Figure 3G). The MTT assay showed that exposure of fibroblasts for 48 h to GW4869 did not induce major cytotoxicity in the cells (Figure 3H).

Consistently, the expression of miR-16, a housekeeping miRNA expressed both by donor cells and exosomes, was significantly reduced in fibroblast-derived exosomes after incubation with GW4869, compared to exosomes derived from untreated cells (1.14 ± 0.03 vs. 0.013 ± 0.0032; p=0.0001), whereas miR-16 endogenous cellular expression was unchanged (1.021 ± 0.293 in GW4869-treated fibroblasts vs. 0.98 ± 0.097 in untreated fibroblasts; p=0.93873) (Figure 3I). All together these data indicate that the release of exosomes from fibroblasts, and the exosomal miRNA content is controlled by nSMase2 and is ceramide-dependent.

### Expression profile of fibrosis-focused miRNAs shows up-regulation of several of them in DMD exosomes

MiRNA isolated from control and DMD exosomes were analysed on a miRNA array containing 84 miRNAs known to play a role in fibrosis. These miRNAs are grouped as: pro-fibrotic, anti-fibrotic, extracellular matrix remodelling and cell adhesion, inflammation, angiogenesis, signal transduction and transcriptional regulation, and epithelial-to-mesenchimal transition clusters. Ct values of miRNAs were analysed using the miScript miRNA PCR Array Data Analysis Tool. Analysis of miRNA profiling showed four significantly up-regulated miRNAs (Figure 4A,B): miR-125b-5p, involved in signal transduction and transcriptional regulation (3.38 ± 0.73; p=0.0051); miR-143-3p, involved in ECM remodelling and cell adhesion (11.14 ± 1.58; p=0.0327); miR-195-5p, involved in angiogenesis and signal transduction and transcriptional regulation (4.42 ± 0.62; p=0.00856) and miR-199a-5p, involved in ECM remodelling and cell adhesion, inflammation and epithelial to mesenchimal transition (11.83 ± 1.23; p=0.0103) (Figure 4C). Real-Time-PCR analysis of control and DMD exosomal RNAs confirmed the increased expression levels of the four miRNAs. In particular, miR-125-5p was 3.18 ± 1.17 (p=0.0005), miR-143-3p was 4.32 ± 1.38 (p=0.00009), miR-195-5p was 1.93 ± 0.69 (p=0.0035) and miR-199a-5p was 2.58 ± 0.99 (p=0.00017) (Figure 4D).

**Figure 4.**
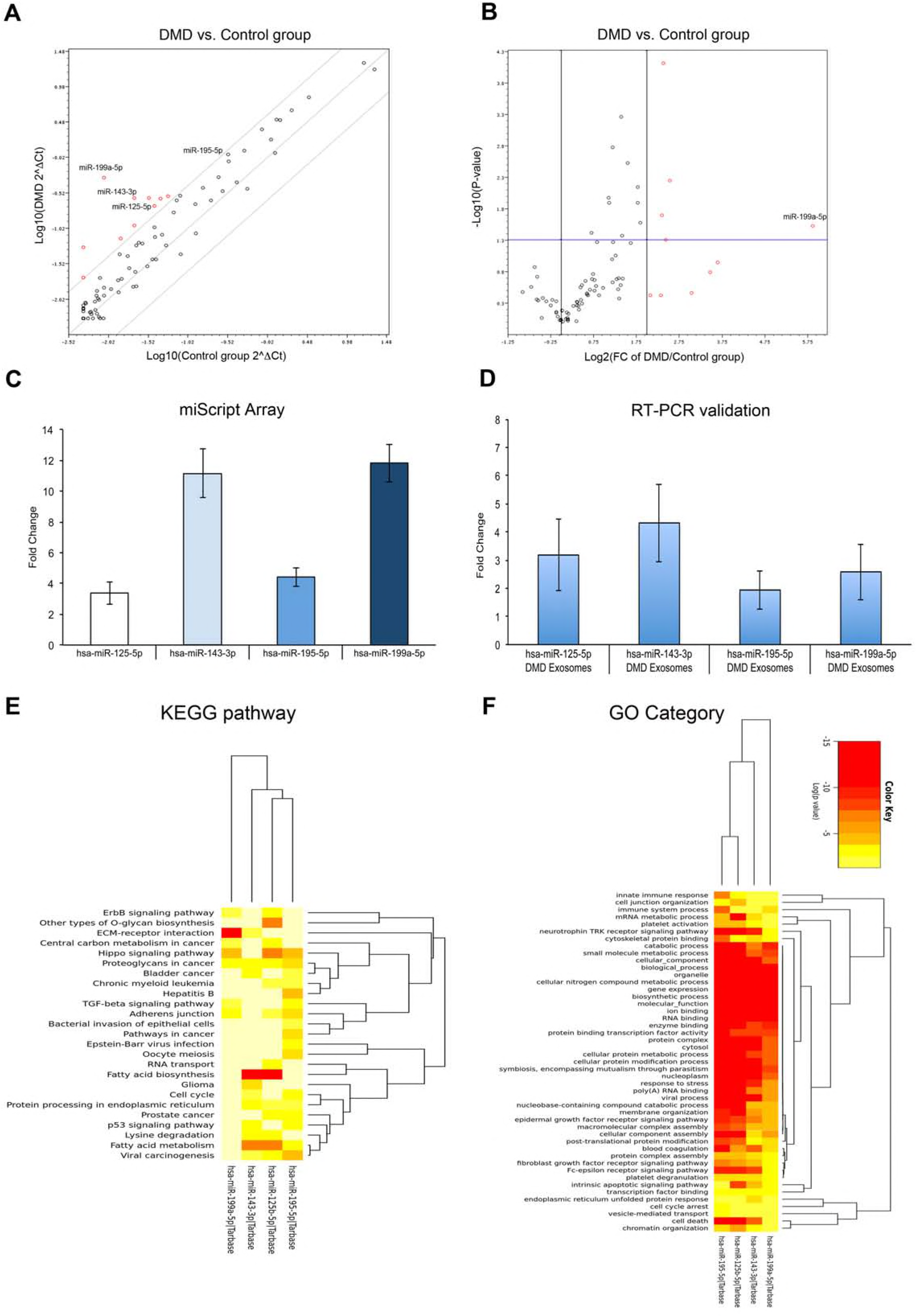
Exosomal miRNA profile analysis. **A, B** Scatter (left) and Volcano (right) plots of miRNA profile expression obtained from comparison of exosomal miRNAs isolated from control and DMD fibroblasts cell lines and analysed on a fibrosis-focused pathway miScript array. **C, D** Fold change of selected miRNAs obtained from miScript array (C) and RT-PCR validation (D) in DMD exosomes. **E, F** Enrichment for the KEGG pathways (E) and GO categories (F) for the selected miRNAs obtained from computational prediction by mirPath v.3 software. Data are presented as mean ± SD, *n* = 3 per group, and analyzed by 2-tailed Student’s t test. P ≤ 0.05.

Computational KEGG and GO predictions (Figure 4E,F) for the four differentially expressed miRNAs identified several target genes involved in important fibrosis-related pathways, including: ECM-receptor interaction (hsa04512; p=2.61E-08), Hippo signalling pathway (hsa04390; p=1.01E-05), adherens junction (hsa04520; p=0.0011729), proteoglycans in cancer (hsa05205; p=0.0010058), Wnt signalling pathway (hsa04310; p=0.0312812), PI3K-Akt signalling pathway (hsa04151; p= 0.040471) and TGF-β1 signalling pathway (hsa04350; p=0.048135). The statistically significant value associated with the identified biological pathways was calculated using mirPath and the Fisher’s exact test.

### Levels of miR-199a-5p are significantly up regulated and transcript and protein levels of its target gene CAV1 are significantly down regulated in DMD fibroblasts, but not in myoblasts

Because of previous studies demonstrating miR-199a-5p involvement in lung (Lino Cardenas *et al*, 2013) and liver (Murakami *et al*, 2011) fibrosis, we focused our attention on this miRNA among those differentially expressed. GO ontology distribution of miR-199a-5p target genes (Figure 5A) indicated that they are involved in a great number of molecular functions, while KEGG analysis showed that several target genes are especially implicated in fibrosis-related pathways (Figure 5B). Furthermore, the STRING software (https://string-db.org) analysis of protein-protein interaction networks (Figure 5C) and of molecular interactions of miR-199a-5p target genes (Figure 5D) pointed to CAV1 as one of the interactors most frequently present in fibrosis-related pathways. CAV1 is in fact one of the experimentally validated target genes of miR-199a-5p (see the conserved sequence seed for CAV1 in Figure 5E) that, besides being involved in various cellular processes, is implicated in the regulation of idiopathic pulmonary fibrosis (Wang *et al*, 2006; Aranda *et al*, 2015).

**Figure 5.**
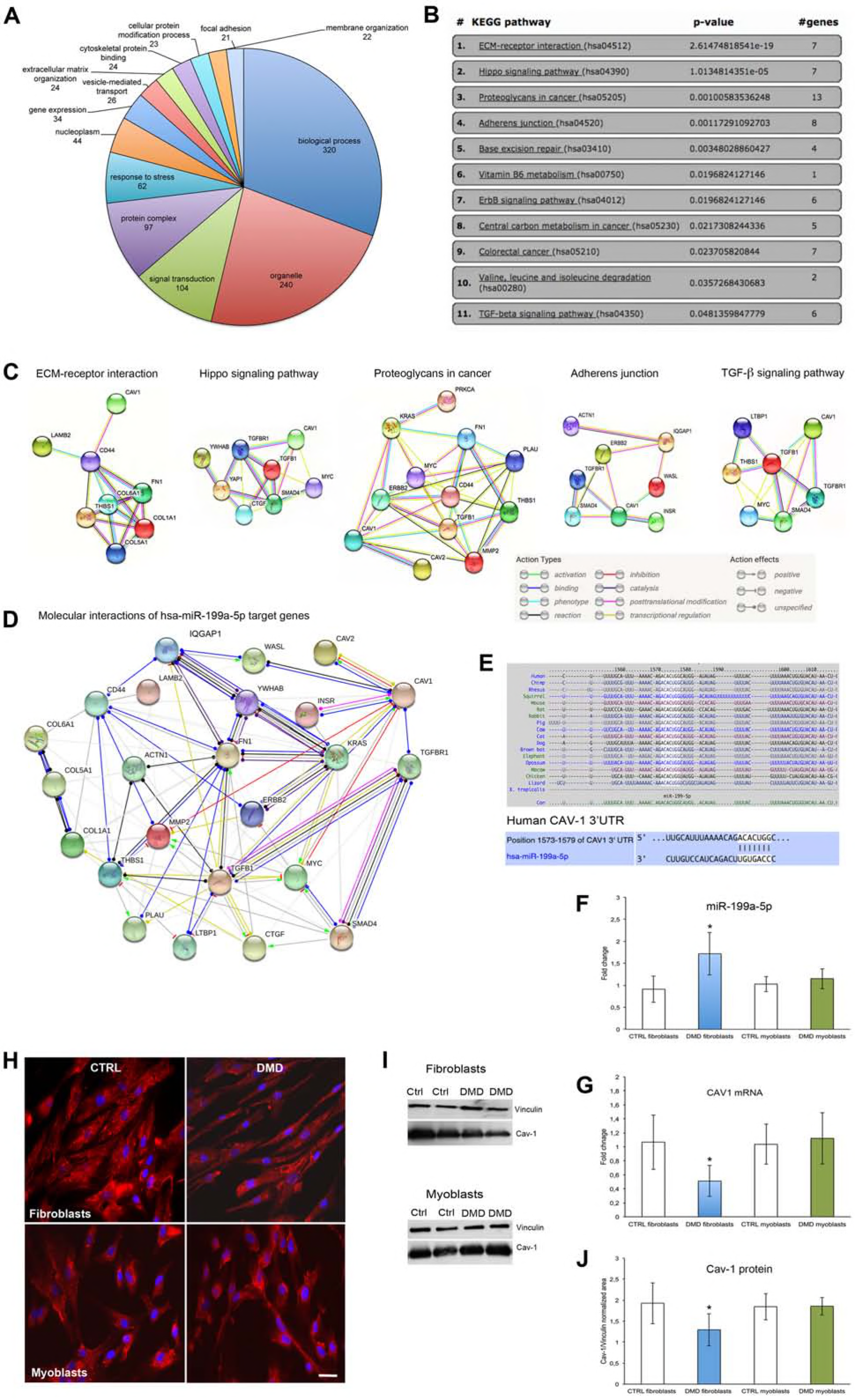
Function and target genes of miR-199a-5p. **A** Distribution and number of miR-199a target genes in different GO categories. **B** KEGG pathway list of miR-199a with related p-value and number of target genes involved in each pathway. **C** String representation of protein-protein interaction of five selected pathways related to fibrosis. **D** String representation of molecular interactions of miR-199a target genes. **E** Phylogenetic conservation of the seed sequence of miR-199a for CAV-1 and related position on the human CAV-1 3’ UTR. **F, G** RT-PCR of miR-199a levels (**F**) and of CAV-1 mRNA expression levels (**G**) in control and DMD fibroblasts and myoblasts. **H** Immunocytochemistry of caveolin-1 in control and DMD fibroblasts and myoblasts. Cell nuclei were stained with DAPI. Scale bar: 10 μm. **I, J** Western blot of caveolin-1 in control and DMD fibroblasts and myoblasts (**I**) and densitometric analysis of western blot bands (**J**). Data are presented as mean ± SD, *n* = 3 per group, and analyzed by 2-tailed Student’s *t* test. P ≤ 0.05.

Levels of miR-199a-5p were significantly increased in DMD fibroblasts compared to control fibroblasts (1.71 ± 0.88 vs. 0.912 ± 0.30; p=0.027) and unchanged in DMD myoblasts (1.15 ± 0.23 vs. 1.027 ± 0.17; p=0.212) (Figure 5F). Accordingly, mRNA levels of the miR-199a-5p target CAV1 were significantly reduced (0.622 ± 0.18 vs. 1.06 ± 0.38; p= 0.009); and protein expression of caveolin-1 was of lower intensity, by immunocytochemistry (Figure 5H), and significantly reduced by densitometric analysis of Western blot bands (Figure 5I,J), in DMD fibroblasts compared to control fibroblasts (1.274 ± 0.36 vs. 1.926 ± 0.48; p= 0.044). CAV1 expression levels were not significantly different in myoblasts both at transcript (0.985 ± 0.28 vs. 1.035 ± 0.28; p= 0.713) and protein level (1.856 ± 0.21 vs. 1.844 ± 0.31; p=0.947) (Figure 5G).

### Horizontal transfer of DMD exosomes to control fibroblasts increases miR-199a-5p and transcript and protein levels of fibrosis-related markers, up-regulates collagen production and activates Akt and ERK pathways

To evaluate a possible pro-fibrotic role of exosomes, control fibroblasts were exposed to DMD exosomes (horizontal transfer). Exposure at different concentrations of DMD exosomes and different timing, significantly increased cell proliferation after 24 h, at the highest exosome concentrations (15 and 30 μg/ml), and after 48 h, at all concentrations tested, compared to untreated fibroblasts (Figure 6A).

**Figure 6.**
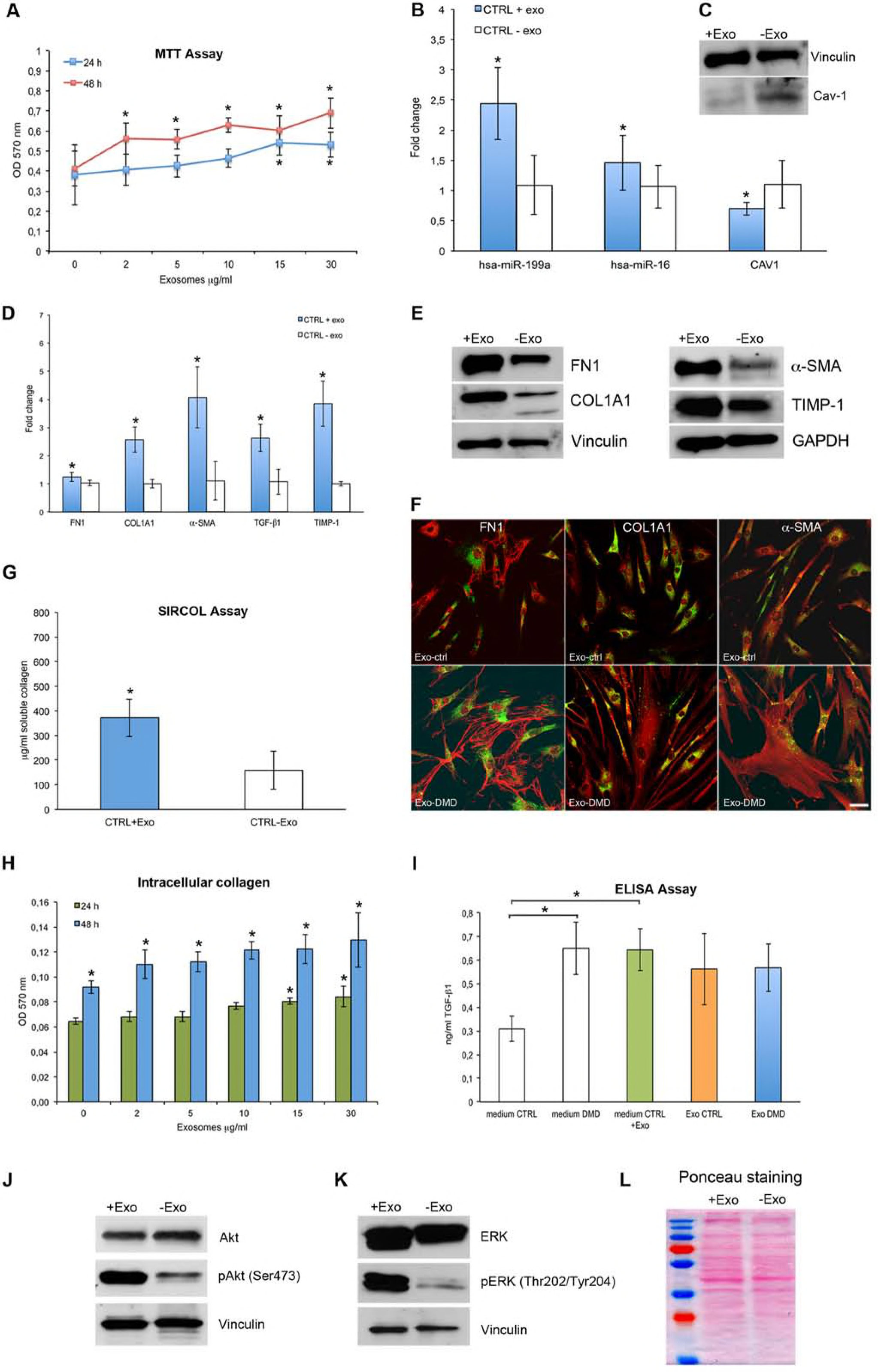
Horizontal transfer of DMD exosomes to control fibroblasts as recipients cells. **A** MTT assay on control fibroblasts after exposure to DMD exosomes for 24 and 48 h. **B** Expression levels of miR-199a, miR-16, and of CAV1 mRNA in control fibroblasts incubated for 48 h with DMD exosomes. **C** Representative western blot of caveolin-1 in control fibroblasts incubated for 48 h with DMD exosomes. **D, E** RT-PCR (**D**) and representative western blots (**E**) of fibrosis-related markers in control fibroblasts after 48 h exposure to DMD exosomes. **F** Immunocytochemistry of FN1, COL1A1 and α-SMA in control fibroblasts incubated for 48 h with PKH67 labelled control exosomes (upper panels) or PKH67 labelled DMD exosomes (lower panels) showing α-SMA positivity in structures resembling stress fibers and myofibroblast-like morphological features of the cells after exposure to DMD exosomes. Scale bar: 10 μm. **G** Sircol assay for quantitation of soluble collagen production in culture media from control fibroblasts before and after incubation for 48 h with DMD fibroblasts exosomes. **H** Quantitation of intracellular collagen deposition in control fibroblasts treated for 24 and 48 h with DMD exosomes. **I** TGF-β1 protein levels, measured by Elisa assay, in the supernatant of control and DMD fibroblasts in basal conditions, in medium of control fibroblasts after incubation with DMD exosomes, and in control and DMD exosomes. **J, K** Representative western blots of Akt (**J**) and (**K**) ERK pathways in control fibroblasts after 48 h of incubation with DMD exosomes. **L** Ponceau S staining as loading control. Data are presented as mean ± SD, n = 3 per group. Unpaired, independent groups of 2 were analyzed by 2-tailed Student’s t test. Multigroup comparisons were analyzed by 1-way ANOVA with Dunnett’s post hoc test. P ≤ 0.05.

Exposure of control fibroblasts to DMD exosomes also significantly increased miR-199a-5p (2.44 ± 0.60 vs. 1.09 ± 0.49; p=0.045) and miR-16 (1.46 ± 0.45 vs. 1.06 ± 0.35; p=0.049) levels, and significantly reduced CAV1 transcript levels (0.8 ± 0.15 vs. 1.10 ± 0.39; p=0.002) (Figure 6B). In addition, a decreased band intensity of the caveolin 1 protein was detected by Western blot (Figure 6C).

Furthermore, transcript levels of the fibrosis-related markers FN1 (1.25 ± 0.16 vs. 1.04 ± 0.104; p=0.001), COL1A1 (2.57 ± 0.74; vs. 1.01 ± 0.15 p=0.001), α-SMA (3.39 ± 1.74 vs. 1.11 ± 0.68; p=0.025), TGF-β1 (2.63 ± 1.48 vs. 1.07 ± 0.44; p=0.007) and TIMP-1 (3.85 ± 0.81 vs. 1 ± 0.08; p= 0.003) were significantly increased after DMD exosome treatment (Figure 6D). An increased intensity of FN1, COL1A1, α-SMA and TIMP-1 protein bands was observed by Western blot in control fibroblasts exposed to DMD exosomes compared to untreated cells (Figure 6E). Staining intensity of FN1, COL1A1 and α-SMA was also increased by immunocytochemistry in control fibroblasts after 48 h exposure to PKH67-labelled DMD exosomes, compared to control fibroblasts exposed to PKH67-labelled control exosomes (Figure 6F). Furthermore, in the fibroblasts treated with DMD exosomes, α-SMA immunostaining appeared positive in structures resembling stress fibers and the cells exhibited morphological features typical of myofibroblasts, not observed in cells exposed to control exosomes (Figure 6F).

Soluble collagen production in recipient control fibroblasts exposed to DMD exosomes was significantly greater than in untreated control fibroblasts (372.36 ± 76.43 μg/ml vs. 159.01 ± 77.55 μg/ml; p=0.00003) (Figure 6G). Intracellular collagen deposition was significantly increased after control fibroblasts were exposed to the highest concentrations of DMD exosomes (15 and 30 μg/ml) for 24 h and to all concentrations tested (5-30 μg/ml) for 48 h (Figure 6H).

Finally, the content of TGF-β1, detected by ELISA, was not significantly different in exosomes derived both from control and DMD fibroblasts (0.56 ± 0.15 ng/ml vs. 0.57 ± 0.10 ng/ml; p= 0.956); however, a significant increase in TGF-β1 levels was observed in culture media of control fibroblasts after 48 h exposure to DMD exosomes, compared to culture media of unexposed cells (0.65 ± 0.11 ng/ml vs. 0.39 ± 0.05 ng/ml, p= 0.001). This value was similar to the value detected in the cell culture media of DMD fibroblasts (0.643 ± 0.088 ng/ml) (Figure 6I).

After exposure of control fibroblasts to DMD exosomes, an increased intensity of the Akt (Ser-473) and ERK phosphorylated bands (p42/44, Thr202/Tyr204) was observed by Western blot, compared to untreated cells (Figure 6K).

### Horizontal transfer of DMD fibroblast-derived exosomes to control myoblasts does not change miR-199a-5p and CAV1 transcript level nor transcript and protein levels of fibrosis-related markers

In control myoblasts exposed to fibroblast-derived DMD exosomes transcript levels of miR-199a (0.89 ± 0.16 vs. 1.03 ± 0.31; p=0.116) and CAV1 (1.12 ±0.36 vs. 1.06 ± 0.44; p=0.852) were not significantly changed compared to untreated cells (Figure 7C); nor were levels of the fibrosis-related markers FN1 (1.09 ± 0.43 vs. 1.02 ± 0.25; p=0.874), COL1A1 (0.83 ± 0.08 vs. 1 ± 0.03; p=0.0984), α-SMA (1 ± 0.03 vs. 1.02 ± 0.2; p=0.892), TGF-β1 (0.84 ± 0.15 vs. 1.09 ± 0.53; p=0.412) and TIMP-1 (0.96 ± 0.04 vs. 1 ± 0.1; p= 0.588) (Figure 7D).

**Figure 7.**
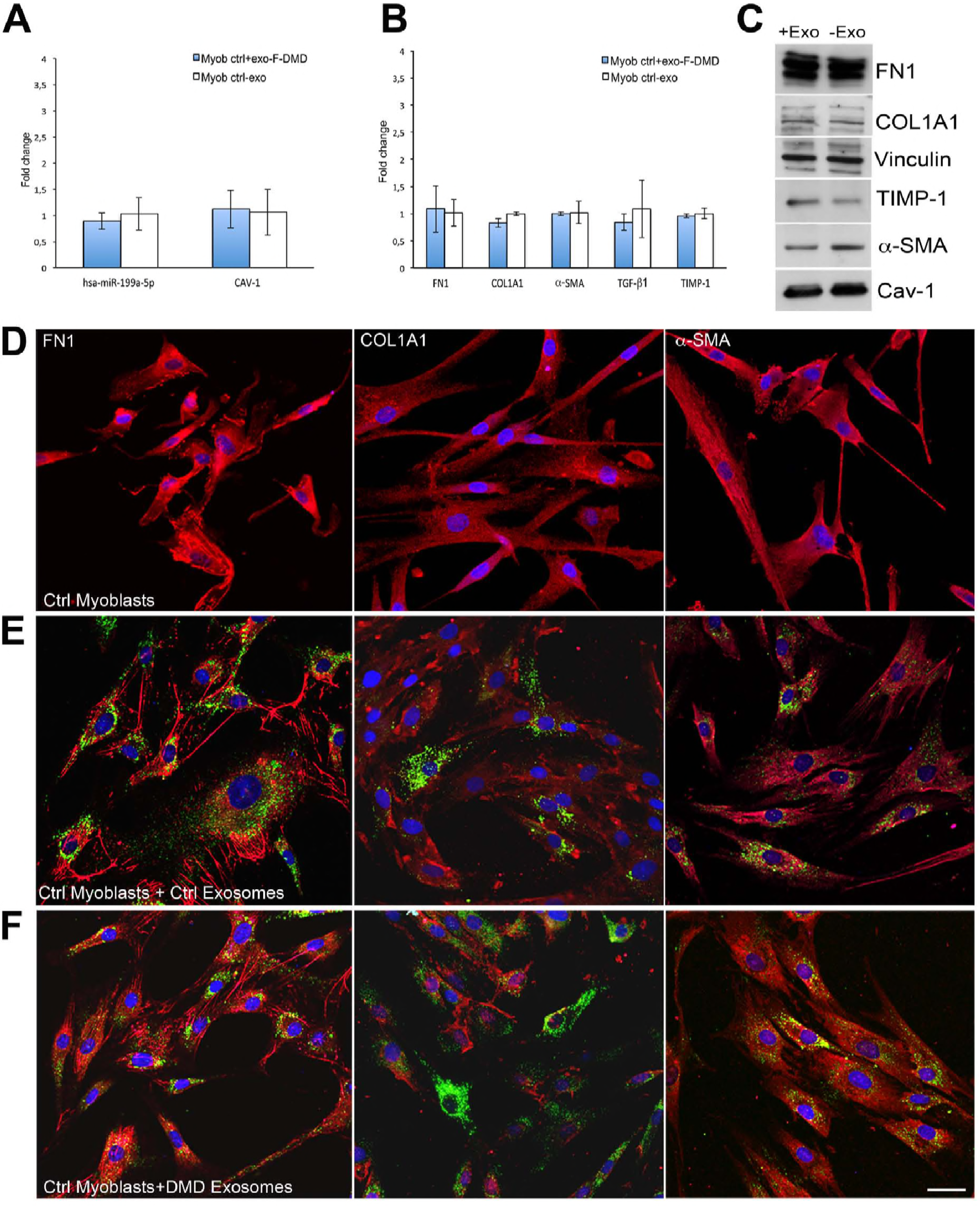
Horizontal transfer of fibroblast-derived exosomes to myoblasts. **A** RT-PCR showing miR-199a levels and CAV-1 transcript levels in control myoblasts after exposure to DMD fibroblast-derived exosomes. **B** RT-PCR of fibrosis related markers after exposure to DMD fibroblast-derived exosomes. **C** Representative western blot of fibrosis related markers in control myoblasts after exposure to DMD fibroblast-derived exosomes for 48 h. **D-F** Immunocytochemistry of FN1, COL1A1 and α-SMA in myoblasts before (**D**) and after incubation for 48 h with PKH67 labelled control (**E**) or DMD (**F**) fibroblast-derived exosomes. Scale bar: 10 μm. Data are presented as mean ± SD, n = 3 per group, and analyzed by 2-tailed Student’s t test. P ≤ 0.05.

No differences in morphology and protein expression were observed by immunocytochemistry and Western blot analysis of myoblasts after incubation with fibroblast-derived DMD exosomes compared to untreated cells and to cells treated with control fibroblast-derived exosomes (Figure 7C-F).

### Transfection of control fibroblasts with miR-199a-5p mimic and inhibitor has opposing effect on miR-199a-5p levels, on fibrosis-related mRNAs and proteins, on collagen production and Akt and ERK pathways

To confirm whether the pro-fibrotic effect of exosomes was to be ascribable at least in part to miR199a-5p, control fibroblasts were transfected with miR-199a-5p mimic and inhibitor.

Transfection efficiency was first evaluated by transfecting the fibroblasts with the miRNA mimic miR-1 and the miRNA inhibitor let-7c, and by measuring their respective targets TWF1 and HMGA2. A significant decrease in TWF1 (ID: Hs00702289_s1) (0.106 ± 0.014; p=0.00108), and a significant increase in HMGA2 (ID: Hs00971725_m1) (24.51 ± 4.77; p=0.00103) mRNA levels were observed compared to untreated fibroblasts (expressed as 1) (Figure 8A).

**Figure 8.**
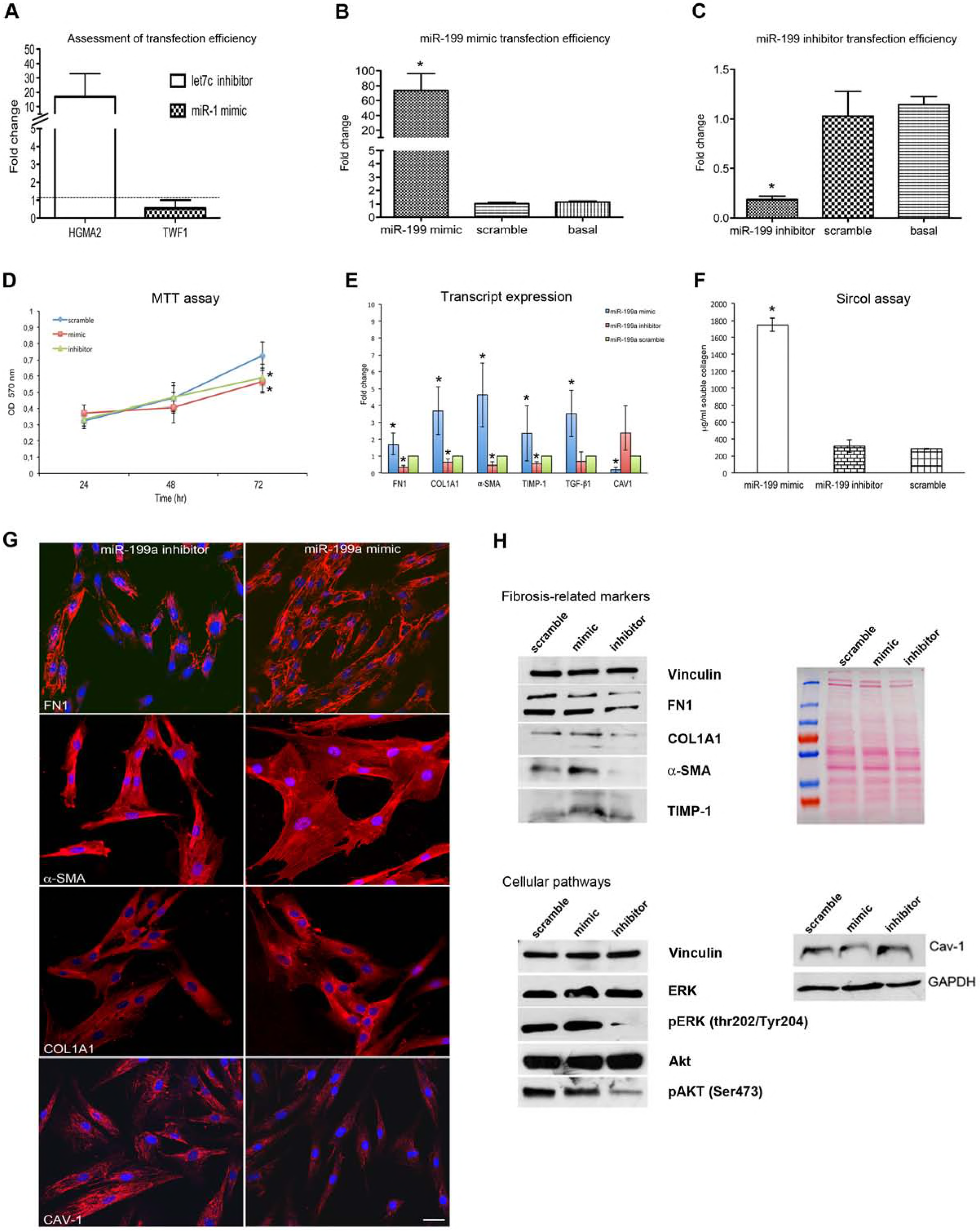
Transfection of control fibroblasts with miR-199a-5p mimic and inhibitor. **A** RT-PCR of HGMA2 and TWF1, respectively target genes of let7c inhibitor and miR-1 mimic, for evaluation of transfection efficiency. **B, C** RT-PCR of miR-199a-5p after transfection with miR-199a-5p mimic (**B**) or with miR-199a-5p inhibitor (**C**) in control fibroblasts. **D** MTT assay performed on control fibroblasts after miR-199a-5p mimic, inhibitor or scramble transfection. **E** RT-PCR of fibrosis related markers in control fibroblasts after transfection with miR-199a-5p mimic, inhibitor or scramble. **F** Sircol assay for quantitation of soluble collagen production in cell medium of control fibroblasts transfected with miR-199a-5p mimic, inhibitor or scramble. **G** Immunocytochemistry of FN1, α-SMA, COL1A1 and caveolin 1 in control fibroblasts transfected with miR-199a-5p inhibitor (left panel) or with miR-199a-5p mimic (right panel). Nuclei were stained with DAPI. Scale bar: 10 μm. **H** Representative western blots of fibrosis related markers, of ERK and Akt cellular pathways and of caveolin-1 in control fibroblasts after transfection with miR-199a-5p mimic, inhibitor and scramble. Data are presented as mean ± SD, n = 3 per group. Unpaired, independent groups of 2 were analyzed by 2-tailed Student’s t test. Multigroup comparisons were analyzed by 1-way ANOVA with Dunnett’s post hoc test. P ≤ 0.05.

Control fibroblasts, similarly transfected with miR-199a-5p mimic, miR-199a-5p inhibitor or scramble, showed, respectively, that levels of miR-199a were significantly increased (73.8 ± 45; p=0.018), significantly reduced (0.22 ± 0.05; p=0.00003), or unchanged (1.04 ± 0.47; p=0.383), compared to basal levels of miR-199a-5p (1.15 ± 0.16) (Figure 8B,C).

Although no significant variations in cell proliferation were observed by MTT assay after 24 and 48 h from transfection with miR-199a-5p mimic or inhibitor, after 72 h, cell proliferation was significantly reduced in fibroblasts transfected with either miR-199a-5p mimic (0.566 ± 0.095; p=0.000002) or inhibitor (0.589 ± 0.085; p=0.00001) compared to scramble transfected cells (0.727 ± 0.06) (Figure 8D).

In control fibroblasts transfected with miR-199a-5p mimic, significant up-regulation of transcript levels of the fibrosis-related genes FN1 (1.70 ± 0.63; p=0.0399), COL1A1 (3.68 ± 1.42; p=0.0307), α-SMA (4.63 ± 1.89; p=0.0294), TIMP-1 (2.33 ± 1.62; p=0.0051) and TGF-β1 (3.53 ± 1.36; p=0.0442); and significant down-regulation of CAV1 transcript levels (0.20 ± 0.16; p= 0.001158) were observed compared to control fibroblasts transfected with scramble miRNA (expressed as 1). On the contrary, after miR-199a-5p inhibitor transfection, significant down-regulation of transcript levels of FN1 (0.35 ± 0.11; p=0.00003), COL1A1 (0.63 ± 0.21; p=0.01204), α-SMA (0.45 ± 0.19; p=0.0085), TIMP-1 (0.53 ± 0.13; p=0.0004), was found, and no significant variation of TGF-β1 (0.67 ± 0.58; p=0.3109) and CAV1 mRNA levels (2.35 ± 1.59; p= 0.0139), compared to control fibroblasts transfected with scramble miRNA (as 1) (Figure 8E).

Soluble collagen production significantly increased in culture media of control fibroblasts transfected with miR-199a-5p mimic (1747.03 ± 76.71 μg/ml) compared to scramble transfected cells (285.64 ± 5.28; p= 0.0001); and no significant variations were observed after transfection with miR-199a-5p inhibitor (316.2 ± 74.11; p= 0.619) (Figure 8F).

By immunocytochemistry, staining intensity of the fibrosis-related markers FN1, COL1A1, and α-SMA appeared increased and staining intensity of caveolin 1 appeared decreased in fibroblasts transfected with miR-199a mimic, compared to cells transfected with miR-199a inhibitor (Figure 8G). Similarly, by Western blot, a markedly increased intensity of the bands corresponding to the fibrosis-related markers FN1, COL1A1, α-SMA and TIMP-1 was observed in fibroblasts transfected with miR-199a-5p mimic, compared to cells transfected with miR-199a inhibitor; vice versa, the caveolin 1 protein band was of reduced intensity after miR-199a mimic transfection and increased after miR-199a inhibitor transfection. Western blot also showed an increase in ERK (p42/44, Thr202/Tyr204) and Akt (Ser473) phosphorylated bands in miR-199a-5p mimic transfected cells and a reduction in miR-199a-5p inhibitor transfected cells (Figure 8H).

### Exosomes loaded with miR-199a-5p mimic or inhibitor induce, respectively, up- or down-regulation of fibrosis-related markers in control fibroblasts

To confirm that the pro-fibrotic effect of DMD exosomes was related to specific miRNA content, we loaded control exosomes with miR-199a-5p. Loading efficiency assessment demonstrated a marked increase of miR-199a-5p mimic levels compared to levels in exosomes loaded with miRNA scramble (23.49 ± 2.23 vs. 1± 0.1; p=0.000302) (Figure 9A).

**Figure 9.**
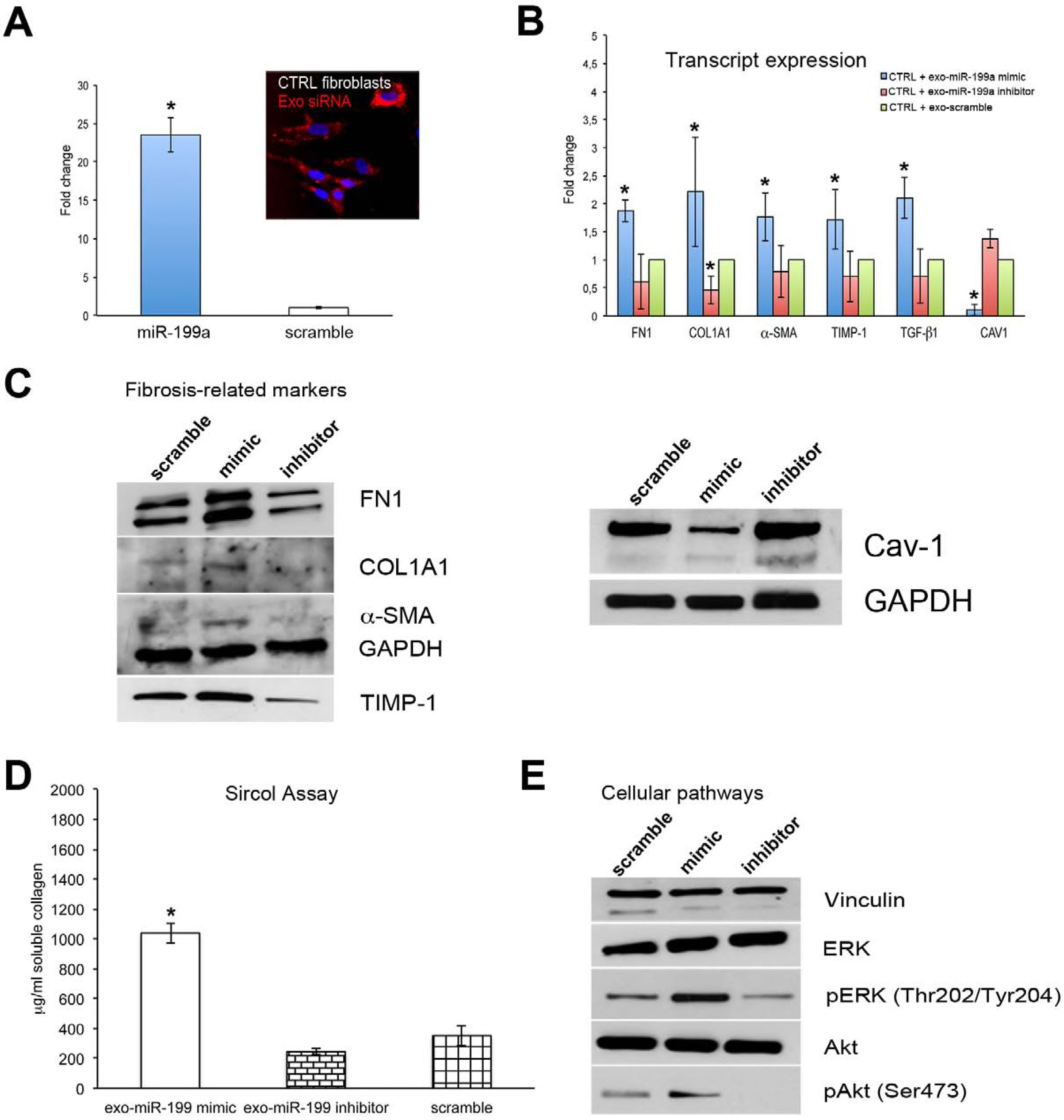
Horizontal transfer of exosomes loaded with miR-199a-5p. **A** RT-PCR for quantitation of miR-199a-5p levels in control exosomes after their loading with miR-199a-5p mimic and scramble. Inset shows transfer of Texas Red fluorescent labelled-siRNA into control exosomes, for evaluation of loading efficiency. Nuclei were stained with DAPI. **B** RT-PCR of fibrosis related markers in control fibroblasts after incubation with exosomes loaded with miR-199a-5p mimic, inhibitor or scramble. **C** Representative western blots of fibrosis related markers and of caveolin-1 in control fibroblasts after incubation with exosomes loaded with miR-199a-5p mimic, inhibitor or scramble. **D** Sircol assay for quantitation of soluble collagen production in cell medium of control fibroblasts after incubation with exosomes loaded with miR-199a-5p mimic, inhibitor or scramble. **E** Representative western blot of ERK and Akt cellular pathways in control fibroblasts after incubation with exosomes loaded with miR-199a mimic, inhibitor or scramble. Data are presented as mean ± SD, n = 3 per group. Unpaired, independent groups of 2 were analyzed by 2-tailed Student’s t test. Multigroup comparisons were analyzed by 1-way ANOVA with Dunnett’s post hoc test. P ≤ 0.05.

When control fibroblasts were incubated with control exosomes loaded with miR-199a-5p mimic, transcript levels of the fibrosis-related genes FN1 (1.74 ± 0.19; p=0.0032), COL1A1 (2.21 ± 0.97; p=0.0497), α-SMA (1.76 ± 0.43, p=0.0302), TIMP-1 (1.71 ± 0.52; p=0.0346) and TGF-β1 (2.10 ± 0.37; p=0.003919) were significantly increased; while CAV1 transcript levels were significantly decreased (0.11 ± 0.09; p= 0.000438), compared with control fibroblasts incubated with exosomes loaded with scramble miR-199a-5p (Figure 9B).

On the contrary, when control fibroblasts were incubated with control exosomes loaded with miR-199a-5p inhibitor, transcript levels of the fibrosis-related genes appeared overall decreased: FN1 (0.60 ± 0.48; p=0.22209), α-SMA (0.79 ± 0.46; p=0.47801), TIMP-1 (0.71 ± 0.45; p=0.332056) and TGF-β1 (0.71 ± 0.47; p=0.352523), but only COL1A1 (0.46 ± 0.25; p=0.015918) levels were significantly reduced; while CAV1 transcript levels were significantly increased (1.37 ± 0.16; p= 0.022), compared with control fibroblasts incubated with exosomes loaded with scramble miR-199a-5p (Figure 9B).

By Western blot a markedly increased intensity of the bands corresponding to the fibrosis-related markers FN1, COL1A1, α-SMA and TIMP-1 was observed in fibroblasts incubated with exosomes loaded with miR-199a-5p mimic, compared to fibroblasts incubated with exosomes loaded with miR-199a-5p inhibitor; and, viceversa, caveolin 1 protein band was of reduced intensity (Figure 9C). Soluble collagen production significantly increased in culture media of control fibroblasts incubated with exosomes loaded with miR-199a-5p mimic (1037.90 ± 66.3 vs. 351.92 ± 17.47; p=0.00478) compared to fibroblasts incubated with exosomes loaded with scramble miR-199a-5p. No significant variations were observed in culture media of control fibroblasts incubated with exosomes loaded with miR-199a-5p inhibitor (244.16 ± 65.15 vs. 351.92 ± 17.47; p=0.1524) (Figure 9D).

Western blot also showed an increase in ERK (p42/44, Thr202/Tyr204) and Akt (Ser473) phosphorylated bands in fibroblasts incubated with exosomes loaded with miR-199a-5p mimic, compared to fibroblasts incubated with exosomes loaded with miR-199a-5p inhibitor (Figure 9E).

### Treatment with DMD exosomes significantly increases muscle fibrosis in mouse tibialis anterior

To further prove the pro-fibrotic role of DMD exosomes, wild type mice were injected into the right tibialis anterior muscle and the contralateral muscle with DMD or control fibroblast-derived exosomes, respectively, after necrosis had been induced by cardiotoxin treatment (Figure 10A). Both groups of muscles exhibited extensive regeneration and areas of increased perimysial and endomysial fibrosis (Supplementary Figure 1). The fibrotic areas, by immunohistochemistry, appeared as collagen I- / collagen VI-positive and contained several fluorescent dots corresponding to exosomes, either single or aggregate. Exosomes were distributed in the endomysium and perimysium and localized in part at the capillaries (Figure 10C and supplementary figure 1).

**Figure 10.**
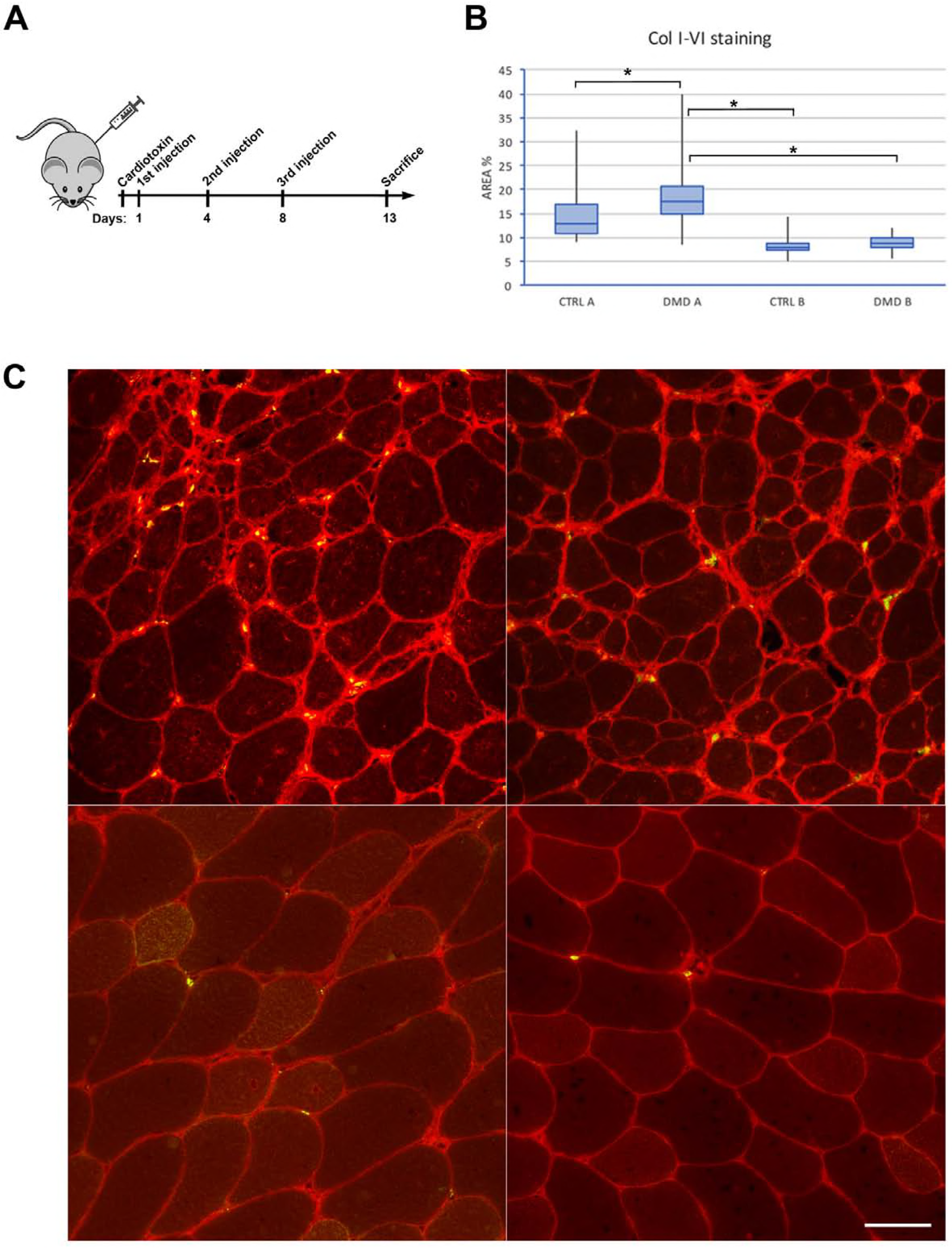
Mouse tibialis anterior treatment with exosomes. **A** Schematic picture of the timeline of mice treatment with DMD and control exosomes after cardiotoxin injury. **B** Box plot of fibrotic areas evaluated as positivity to collagen I-VI staining. **C** Immunohistochemistry of mouse tibialis anterior after treatment with DMD exosomes (left panel) or control exosomes (right panel) showing more marked fibrosis in areas with highest number of exosomes (green dots). Scale bar: 100 μm. Data distribution is presented as quartiles, maximum and minimum values, and medians, n = 32 per group. The linear regression with interacted dummy variables was employed to assess differences among groups. P ≤ 0.05.

Assessment of fibrosis in muscles by comparing the areas of collagen I/collagen VI positivity, revealed that muscles treated with DMD exosomes and containing more than 10 dots had significantly more fibrosis than those injected with control exosomes. Also, muscles treated with DMD exosomes and containing more than 10 dots had significantly increased fibrosis (mean value: 17.90) compared to all other groups. In particular, the fibrotic mean area in muscles treated with DMD exosomes and containing less than 10 dots (DMD B) was 8.73 (p<0.001); in muscles treated with control exosomes and containing more than 10 dots (control A) mean area was 15.46 (p=0.038); and in muscles treated with control exosomes containing less than 10 dots (Control B) mean area was 8.08 (p<0.001) (Figure 10B).

## DISCUSSION

We have investigated for the first time the possible role of DMD fibroblast-derived exosomes in inducing trans differentiation of normal fibroblasts to myofibroblasts and the contribution of exosomes to the modulation of muscle fibrosis. Phenotypic modifications induced by horizontal transfer of exosomal miRNAs to recipient cells are well documented in cancer transformation (Zhang *et al*, 2015; Subramanian *et al*, 2016) where tumour-derived extracellular vesicles (EVs) can induce noncancerous cells to become cancerous. Horizontal transfer of exosomal content is also implicated in physiological processes, such as immune system regulation (Robbins & Morelli, 2014) and nervous system development and homeostasis (Frühbeis *et al*, 2013). Both in pathological and physiological conditions the transfer of information by EVs between cells, is carried out by mRNAs and various forms of non-coding RNA, such as miRNAs and long non coding RNAs (Ogorevc *et al*, 2013; Lopatina *et al*, 2016).

In the present study we first demonstrated that exosomes are indeed produced and released by fibroblasts and myoblasts isolated from DMD and control muscle biopsies. Then we have shown that exosomes can be up taken by fibroblasts in a time and temperature dependent manner without cell toxicity. We have also found that exosome release and uptake similarly occurs in myoblasts (data not shown).

Mechanisms by which exosomes and EVs in general can enter cells and deliver their cargo include phagocytosis, macropinocytosis, clathrin-mediated or fibronectin-mediated endocytosis, and plasma or endosomal membrane fusion. (Mulcahy *et al*, 2014). Biogenesis and release of exosomes can occur either via the ESCRT machinery, or ceramide (nSmase2) pathway, or both (Chen *et al*, 2017). We have demonstrated that exosome uptake requires an intact cytoskeleton, as verified by cytochalasin B treatment, and occurs without involvement of the EDA domain of fibronectin. Exosome partial co-localization with endosomes and lysosomes, but not with endoplasmic reticulum, suggests that, after uptake and internalization, exosomes can diffuse in the cytoplasm. The rapid internalization of exosomes into the cells suggests that it may take place via phagocytosis or macropinocytosis rather than via plasma membrane fusion. We have also shown that the release of exosomes and exosomal miRNA content from fibroblasts occurs via a ceramide-dependent pathway, although we cannot exclude that an ESCRT pathway could also be involved. Then, to further characterize DMD fibroblast-derived exosomes as carriers of miRNAs we analysed, by a focused pathway array related to fibrosis, their miRNA profiling. By computational analysis we identified four miRNAs significantly up-regulated in DMD fibroblast-derived exosomes compared to control exosomes. Of note, expression of the same miRNAs did not differ between control and DMD myoblast-derived exosomes. Among the altered miRNAs, we focused our attention on miR-199a-5p for its known involvement in myofibroblast differentiation and in lung and liver fibrosis (Lino Cardenas *et al*, 2013; Murakami *et al*, 2011). Lino Cardenas *et al.* (2013), in particular, demonstrated that silencing of miR-199a-5p in MRC-5 cells, strongly inhibited TGF-β1-mediated differentiation of fibroblasts into myofibroblasts, wound healing and SMAD signalling and suggested that the fibromiR miR-199a-5p promotes pathogenic activation of fibroblasts in response to TGF-β1 by regulating CAV1 as target gene.

We found that, compared to control cells, miR-199a-5p was significantly increased in DMD fibroblasts, but not in DMD myoblasts. These data confirm, once more, our previous findings that fibroblasts from DMD patients have pro-fibrotic features and indicate that these features are related at least in part to their exosomal content. As consequence of increased miR199a-5p expression, its target gene CAV1 was reduced in DMD fibroblasts, but not in DMD myoblasts. MiR-199a-5p target genes are involved in a great number of molecular functions, as also indicated by GO analysis, and several of them are especially implicated in fibrosis-related pathways. Among them, CAV1 is involved in fibrosis as negative modulator of TGF-β1-induced ECM production and, consequently, of SMAD signalling (Wang *et al*, 2006; Le Saux *et al*, 2008).

Third, we observed an increase in miR-199a-5p and concomitant reduction in CAV1 expression levels in addition to a series of fibrosis-related events that included increase in fibrosis-related markers, transdifferentiation of fibroblasts to myofibroblasts, increased production of soluble collagen and increased deposition of intracellular collagen, when DMD fibroblast-derived, but not myoblast-derived, exosomes were transferred to recipient cells. Transcript and protein levels of TGF-β1 were also significantly increased in recipient cells, although TGF-β1 exosomal content was not significantly different in control and DMD exosomes. A likely explanation is that the increased levels of TGF-β1 is not due to a direct horizontal transfer of TGF-β1 by DMD exosomes, but due to the downstream effect of reduced levels of CAV1, target gene of the horizontally transferred miR-199a-5p. Caveolin-1 in fact, as principal component of caveolae, is involved in the internalization and proteasomal degradation of the TGF-β receptors through recruiting SMURF/SMAD7 TGF-β-inhibitory complex, a process that reduces TGF-β1 signalling and potentially restores normal homeostasis of ECM (Di Guglielmo *et al*, 2003). In addition, caveolin-1 exerts its anti-fibrotic role through other mechanisms pivotal for myofibroblast differentiation such as cell adhesion (Grande-Garcia & del Pozo, 2008), mechanotransduction (Parat, 2009) and cell proliferation (Cho *et al*, 2004). Down-regulation of caveolin-1 expression has been found in different fibrotic conditions, including in human fibroblasts during atrial fibrillation (Yi *et al*, 2014), in patient lung and skin with systemic sclerosis (Del Galdo *et al*, 2008) and in bronchiolar epithelium in lung fibrosis (Odajima *et al*, 2007). Furthermore, down-regulation of caveolin-1 expression was found to correlate with pathological activation of PI3K/Akt in idiopathic pulmonary fibrosis (Xia *et al*, 2010), and to induce activation of the signalling molecules ERK, JNK and Akt and consequent over-expression of collagens and α-SMA (Wang & Zang, 2006). To this regard, we indeed found activation of the PI3k/Akt and ERK/MAPK pathways after horizontal transfer of DMD exosomes to control fibroblasts.

Fourth, to prove that the exosome-induced pro-fibrotic effect was to be related specifically to the transfer of the exosomal fibromiR, miR-199a-5p, we devised two experiments, one by transfection with miR-199a-5p mimic, inhibitor, or scramble, and one by loading control exosomes with miR199a-5p mimic, inhibitor, or scramble. Both experiments indeed showed opposing effects on fibrosis-related transcripts and proteins, on collagen production and on Akt and ERK pathways, supporting what we observed with the horizontal transfer experiment.

Finally, a functional evidence of the role of exosomes in muscle fibrosis was obtained by injecting exosomes derived from DMD and control fibroblasts into the tibialis anterior of mice pre treated with cardiotoxin to induce muscle necrosis and subsequent regeneration. Extensive regeneration and perimysial and endomysial increase in collagen expression was observed with both treatments; however, DMD exosomes induced a significantly greater ECM production compared to control exosomes.

In conclusion, our experiments have shown that the cross talk between DMD exosomes and control fibroblasts, as recipient cells, induced the horizontal transfer of miR-199a-5p and the consequent reduction of caveolin-1 expression, suggesting miR-199a-5p and caveolin-1 as potential therapeutic targets in muscle fibrosis. The most likely consequence of such exosomal horizontal transfer is a rapid induction of Akt and Erk phosphorylation, which in turn favours the expression of α-SMA, the synthesis of collagen I and fibronectin and a further activation of TGF-β1 signalling, finally inducing the trans-differentiation of control fibroblasts to myofibroblasts. In DMD muscle, exosomes produced by residential pro-fibrotic fibroblasts may thus contribute to maintain the activated status of myofibroblasts favouring the perpetuation of fibrosis.

As fibrotic conditions remain a major burden to the public health system, novel therapies that target miRNAs may potentially be promising in the clinic. Furthermore, exosomes, in addition to be carrier of harmful messengers to the tissues, could also be charged with positive anti-fibrotic molecules, either proteins, mRNAs, or miRNAs, thus representing a powerful tool to modify fibrotic tissue degeneration.

## Materials and Methods

### Reagents

Desmin monoclonal antibody (mAb) (M0760), prolyl-4-hydroxylase mAb (M0877), and HSP70 polyclonal antibody (pAb) (A0500) were purchased from Dako. CD63 (TS63) mAb, and EEA1 (PA1-063A) pAb were from Thermo Scientific. Alix (1A12) mAb, fibronectin FN1-IST-9 (SC-59826) and FN1-EP5 (SC-8422) mAbs, caveolin-1 (SC-894) pAb, and collagen VI (SC-20649) pAb were from Santa Cruz Biotechnology. LAMP-1 (H4A3) mAb was from DHSB Developmental Studies Hybridoma Bank. GM130 (610823) mAb was from BD Transduction Laboratories. Calnexin (AB2301) pAb, α-SMA (1A4) mAb, vinculin (V4505) mAb, and collagen 1 (C2456) mAb were from Sigma-Aldrich. Protein kinase B (Akt) (9272) pAb, phosphorylated Akt (pAkt) (9271) pAb, Erk extracellular signal-regulated kinase (Erk) (9102) pAb, and phosphorylated Erk (pErk) (9101) pAb were from Cell Signaling Technology. Tissue inhibitor of metalloproteinase-1 (TIMP-1) (MAB-12070) mAb was from Immunological Sciences. GAPDH (MAB374) mAb was from Millipore Merck. α-tubulin (TU-01) mAb was from GE Healthcare Bio-Sciences. Collagen I (AB765P) pAb was from Chemicon. Biotin-conjugated secondary antibodies (GAM 115-065-062; GAR 111-065-144) and peroxidase-conjugated streptavidin (016-030-084) for Western blot were from Jackson ImmunoResearch Laboratories. ECL (RPN2209) chemiluminescence reagent was from GE Healthcare Bio-Sciences. Cytochalasin B (C6762), PKH67 Green Fluorescent Cell Linker Mini Kit (PKH67GL), GW4869 (D1692) and RGD peptide (8052) were from Sigma-Aldrich. Total Exosome Isolation kit (4478359), CellTracker Blue CMAC (C2110), and total Exosome RNA & Protein Isolation Kit (4478359) were from ThermoFisher. miScript II RT kit (218161) Sciences and miScript SYBR Green PCR Kit (218073) were from Qiagen. miScript miRNA PCR Array human fibrosis (MIHS-117Z) was from SABiosciences, Qiagen. Exo-Fect Exosome Transfection Kit (EXFT10A-1) was from System Biosciences SBI. Sircol assay (S1000) was from Biocolor.

### Human muscle cell cultures

Cell cultures were provided by the Italian Telethon Network of Genetic Biobanks http://biobanknetwork.telethon.it/Pages/View/Documents (Mora et al. 2017). All samples obtained from the biobank had written informed consent using a form approved by the local Ethics Committee. Research was conducted according to protocols approved by the Carlo Besta institutional review board.

Fibroblasts and myoblasts were obtained by immunomagnetic selection, as described in (Zanotti *et al*, 2010), from primary cell cultures derived from muscle biopsies of three DMD patients (aged 3-6 years) and three age-matched controls (age 2-6 years). Fibroblasts were cultured in DMEM (Lonza Group Ltd), supplemented with 10% heat-inactivated fetal bovine serum (FBS) (Gibco Life Technologies), 1% penicillin−streptomycin (Lonza), and 2 mM L-glutamine (Lonza), at 37°C in 5% CO_2_. Immunocytochemistry using prolyl-4-hydroxylase antibody to assess purity of fibroblast populations showed that 100% cells were positive.

Myoblasts were cultured in DMEM supplemented with 20% FBS, 1% penicillin−streptomycin, 2 mM L-glutamine, 10 μg/ml insulin (Sigma-Aldrich), 2.5 ng/ml basic fibroblast growth factor (Gibco), and 10 ng/ml epidermal growth factor (Gibco). Immunocytochemistry using desmin and counting nuclei of desmin-positive cells over total nuclei, to assess purity of myoblast populations, showed that 93.5 % of control myoblasts and 94.02 % of DMD myoblasts were desmin-positive.

Experiments were performed on early passages (3−10 passages) of normal and dystrophic fibroblasts and myoblasts. Both cell types were subjected to the mycoplasma test using the MycoAlert Plus mycoplasma detections kit (Lonza).

### Exosome isolation

Control and DMD fibroblasts or myoblasts at 70% of confluence were incubated in DMEM containing 10% exosome-depleted FBS (Gibco) for 48 h. Subsequently, the media from control and DMD fibroblasts or myoblasts were collected and centrifuged at 2,000 × g for 30 min at 4°C to thoroughly remove cellular debris. The supernatants were filtered through a 0.22-μm filter (Millipore, Billerica, MA) to remove microvesicles. Exosomes were isolated using Total Exosome Isolation kit (4478359) according to manufacturer’s specifications. Briefly, the supernatant was mixed with Total Exosome Reagent and incubated for 24 h at 4 °C. Samples were centrifuged at 10,500 × g for 2 h (Sigma 3-18K centrifuge with 19777-H rotor) and the supernatant removed. The exosome pellets were resuspended in 100 μl of PBS and stored at −20°C for protein or RNA extraction.

### Protein isolation and Western blot

Proteins from cells and exosomes were extracted in 100 μl RIPA buffer (10 mM Tris−HCl pH 7.4, 150 mM NaCl, 1 mM EDTA, 1 mM EGTA, 0.5% Nonidet-P, 1% Triton X-100) containing a protease inhibitor cocktail (P8340) (Sigma-Aldrich). Exosomal and cellular proteins were quantified with the DC Protein Assay kit (Pierce Biotecnology). For Western blot, 5 μg of exosomal or cellular proteins were mixed to an equal volume of 2X Laemmli buffer (0.5 M Tris−HCl, pH 6.8, 20% glycerol, 2% SDS, 5% 2-mercaptoethanol, and 1% bromophenol blue), boiled for 5 min and loaded onto 10% or 12.5% SDS polyacrylamide gel, electrophoresed and transferred to nitrocellulose membranes (Schleicher and Schuell,). After Ponceau S (Biorad) staining, membranes were probed with primary antibodies to: CD63, Alix, HSP70, LAMP-1, GM130, or calnexin.

Extracts of recipient cells obtained after exposure to DMD exosomes were tested with the following antibodies: FN1-EP5, collagen 1, α-SMA, TIMP-1, protein kinase B (Akt), phosphorylated Akt (pAkt), Erk extracellular signal-regulated kinase (Erk), phosphorylated Erk (pErk), or caveolin 1. Vinculin or GAPDH were used as internal loading control. Biotin-conjugated appropriate secondary antibodies were then applied, followed by peroxidase-conjugated streptavidin, and detection with the ECL chemiluminescence reagent.

To compare protein patterns from isolated exosomes and whole cell lysates, PageBlue Protein staining (ThermoFisher) was used to stain electrophoretic gels according to manufacturer’s instructions.

### Electron microscopy

Exosome pellets were resuspended in 2% paraformaldehyde in phosphate buffer for 30 min, and washed in phosphate buffer. The fixed samples were absorbed onto formvar-coated copper grids for 20 min in a dry environment and fixed in 2 % glutaraldehyde for further 5 min. After being rinsed in distilled water, samples were stained with Uranyl Acetate Replacement Stain (Electron Microscopy Sciences) for 1 min. Excess liquid was removed from the grid using filter paper, and grids were stored at room temperature until imaging. Imaging was done in a FEI Tecnai electron microscope (FEI).

### Exosome uptake

For exosome uptake experiments, freshly isolated fibroblast- or myoblast-derived exosomes were labelled with the PKH67 Green Fluorescent Cell Linker Mini Kit according to the manufacturer’s protocol. Briefly, exosomes diluted in PBS were added to 1 ml Diluent C. In parallel, 4 μl PKH67 dye were added to 1 ml Diluent C and incubated with the exosome solution for 4 min. Labelling was stopped by addition of 2 ml 0.5% BSA/PBS. The labelled exosomes were centrifuged at 10,500 g for 2 h, and the pellet obtained was diluted in 100 μl PBS. To remove the excess of dye, the PKH67-labelled exosomes were purified on Exosome Spin Columns MW 3000 (Invitrogen) according to the manufacturer’s instructions. PKH67-labelled exosomes were added to the culture medium of recipient cells growing onto microscope coverglass slides and incubated at 37°C, for 30 min, or 2, 4 or 24 h. As control of active exosome up-take, control fibroblasts or myoblasts were also incubated with PKH67-labelled exosomes for 4 h at 4°C and α-tubulin used as cytoskeletal marker.

### Cytochalasin B treatment

To evaluate the role of actin in exosome internalization, exosomes were added to cells (growing onto microscope slides) either treated or untreated with cytochalasin B, 10 μg/ml for 4 h. The cells were then washed three times with PBS, fixed in 4% paraformaldehyde for 15 min at room temperature, washed again three times with PBS, and the nuclei counterstained with DAPI (Sigma-Aldrich) or TO-PRO-3 (Invitrogen). Slides were mounted with Mowiol antifading reagent (Invitrogen) and image acquisition was performed with a Leica confocal TCS SP8 multiphoton microscope equipped with hybrid and argon lasers using DIC and fluorescence acquisition (Leica Microsystems). The amount of PKH67-labelled exosomes up-taken by fibroblasts was evaluated by counting the exosomes in the cytoplasm of fibroblasts untreated or treated with cytochalasin B. As cytotoxicity control, cells were incubated with cytochalasin B and stained with vital CellTracker Blue CMAC (25 μM for 45 min at 37°C).

### Immunocytochemistry

After exposing for 4 h control fibroblasts to DMD exosomes labelled with PKH67, co-localization with the antibodies EEA-1 (early endosome marker), LAMP-1 (lysosome marker), calnexin (endoplasmic reticulum marker), or with LysoTracker Red DND-99 (50 nM, ThermoFisher), was assessed with the Leica confocal microscope. Each picture consisted of a z-series of an average of 10 images of 1024 × 1024 pixel resolution with a pinhole of 1.0 airy unit. Fluorescence emission was collected by a HC PL APO CS2 40×/1.10 water-immersion or 63×/1.40 oil-immersion objective. Co-localization was performed using the open source Image J Fiji software version 2.0.0 (*https://imaaei.nih.Qov/iy*).

### RGD peptide treatment

To assess whether exosome uptake was mediated by fibronectin, DMD fibroblasts were treated with the RGD peptide (100 μM), a cell adhesion motif present within fibronectin. Fibroblast culture medium was collected and processed for exosome isolation as described above. Isolated exosomes were then labelled with PKH67 and added to control fibroblasts plated on coverglass slides for 4 h. Cells were extensively washed with PBS, fixed with 4% paraformaldehyde, and nuclei were counterstained with DAPI. The up-take by control fibroblasts of exosomes derived from RGD pre-treated or untreated DMD fibroblasts was evaluated by counting PKH67-labelled exosomes in the cell cytoplasm as before. Western blot analysis was carried out on control and DMD exosomes and cell extracts to evaluate expression of the alternative spliced domain of FN1 (EDA-FN1) using the FN1-IST-9 antibody.

### Treatment of fibroblasts with nSmase2 inhibitor (GW4869)

Fibroblasts were treated with 10 μM nSmase2 ceramide biosynthesis inhibitor (GW4869, Sigma) in DMEM containing 10% exosome-depleted FBS for 48 h. As control, cells were treated with DMSO at the same concentration as the inhibitor. After incubation, medium was collected for exosome purification and cells were collected for miRNA isolation. The expression of miR-16 (Assay ID 000391-miRBase Accession Number: MIMAT0000069), a miRNA expressed in both donor cells and exosomes, was determined.

### MTT assay

The MTT assay was employed to assess cell proliferation (1) after exposure of cells to exosomes, (2) after transfection with hsa-miR-199a mimic, inhibitor or scramble of control fibroblasts, and (3) after exposure to GW4869 nSmase2 inhibitor.

For all 3 tests control fibroblasts were seeded at 2 × 10^4^ cells/well density in 96-well plates in DMEM supplemented with 10% FBS.

For test (1), after overnight plating, cells were washed three times with PBS and incubated with exosomes (1 μg/ml, 10 μg/ml, and 25 μg/ml in DMEM containing 10% exosome-depleted FBS) for 48 h at 37°C.

For test (2), after overnight plating, cells were transfected with 15 pmol of miR-199a mimic, inhibitor or scramble, and cell viability assessed at 24, 48, 72 and 96 h.

For test (3), after overnight plating, cells were washed three times with PBS and GW4869 was added to cells (0, 0.5, 1, 2.5, 5, 7.5, 10 and 20 μM in DMEM containing 10% exosome-depleted FBS) for 24 h.

In all three experiments at the end of incubation, 20 μl of 5 mg/ml MTT (3- [4.5-Dimethylthiazol-2-yl]-2.5-diphenyltetrazolium bromide, Sigma-Aldrich) were added to each well and plates were incubated for 3 h at 37°C. After washing with PBS, 150 μl of MTT solvent solution (4 mM HCl, and 0.1% Nonidet P-40 in isopropanol) were added to each well. Absorbance was read, after 15 min, at 570 nm on a Victor Wallac 1420 multi-label reader (Perkin-Elmer).

### RNA extraction from exosomes

RNA was extracted from exosomes using Total Exosome RNA & Protein Isolation Kit according to the manufacturer’s protocol. RNA was finally eluted in 100 μl of nuclease-free water. Quantification and quality check of extracted RNA was performed with Qubit microRNA Assay kit (Life Technologies) according to manufacturer’s instructions.

### Exosomal RNA reverse transcription for miRNA profiling

Exosomal RNA was reverse transcribed using miScript II RT kit according to the manufacturer’s protocol. In brief, 150 ng RNA were incubated at 37°C for 1 h in 5x miScript HiSpec buffer, 10x miScript Nucleics Mix and miScript Reverse Transcriptase Mix in 20 μl final volume. After inactivation at 95°C for 5 min, cDNA was diluted with 200 μl RNase-free water and used for miRNA PCR array.

MiRNA PCR array was performed using the miScript SYBR Green PCR Kit on custom printed 96-well miScript miRNA PCR Array human fibrosis. A set of controls, enabling data analysis, assessment of reverse transcription performance, and assessment of PCR performance, is included on each plate. The miScript miRNA real-time PCR was performed using ABI Prism 7000 (Applied Biosystem) with an initial activation step of 95°C for 15 min, followed by 40 cycles of 3-step cycling (denaturation, 15 sec, 94°C; annealing, 30 sec, 55°C; and extension, 30 sec, 70°C). Ct values were exported and analysed using the miScript miRNA PCR Array Data Analysis Tool (http://pcrdataanalysis.sabiosciences.com/mirna, SABiosciences, Qiagen). The relative expression was calculated using the ΔΔCT method [(relative gene expression = 2-(ΔCT sample-ΔCT control)] and is presented as fold increase relative to control levels expressed as 1. Ct values were normalized versus housekeeping genes (*SNORD61*, *SNORD68*, *SNORD95*, *SNORD96A* and *RNU6B*) and the Ct cut-off applied was 35.

Computational predictions of target genes were performed using the miRecords (http://c1.accurascience.com/miRecords/), TargetScan (http://www.targetscan.org/) and PicTar (http://pictar.mdc-berlin.de/) databases and prediction tools available online. To further elucidate the biological processes and corresponding metabolic networks regulated by the identified miRNAs, pathways were identified with mirPath v.3 (http://snf-515788.vm.okeanos.grnet.gr/) by performing Kyoto Encyclopedia of Genes and Genomes (KEGG) and gene ontology (GO) pathway analyses (Vlachos *et al*, 2015).

### TaqMan miRNA assay for miRNA array validation

For qPCR confirmation, total RNA was isolated from exosomes as described above. Four differentially expressed miRNAs: miR-199a-5p (Assay ID 000498, miRBase Accession Number: MIMAT0000231), miR-125b-5p (Assay ID 000449, miRBase Accession Number: MIMAT0000423), miR-195-5p (Assay ID 000494, miRBase Accession Number: MIMAT0000461), and miR-143-3p (Assay ID 002249, miRBase Accession Number: MIMAT0000435), were further quantitated by TaqMan miRNA assays (Applied Biosystems). The expression of each miRNA was normalized to the expression of *RNU6B* (Assay ID 001093) and determined as the ratio of the target miRNA to the *RNU6B* calculated by 2−ΔΔCt, where ΔΔCt = CtTarget − CtU6B. Quantitative PCR data were expressed as “fold change” relative to control levels expressed as 1. Samples were processed in triplicate.

### Incubation of recipient cells with exosomes

To verify the biological activity of exosomes, recipient cells (control fibroblasts or myoblasts) were incubated with exosomes isolated from DMD fibroblasts or myoblasts. Incubation was for 48 h in DMEM containing 10% exosome-depleted FBS plus equal amounts of freshly isolated exosomes. At the end of incubation, cells and cell culture media were collected, respectively, for protein and RNA extraction and for Sircol assay.

### Transcript quantitation by RT-PCR in recipient cells

Total RNA was isolated from fibroblasts or myoblasts using TRI Reagent (Ambion) according to manufacturer’s instructions, and checked spectrophotometrically for quantity and purity, using a Nanodrop 2000C spectrophotometer (Thermo Scientific). RNA aliquots (1 μg) were reverse transcribed in the presence of 5x first strand buffer (Life Technologies), 1 mM each deoxynucleoside triphosphate, 8 pM random hexamers, 10 μM dithiothreitol, 1 IU/μl RNAse inhibitor (Roche Molecular Biochemicals), and 10 IU/μl M-MLV reverse transcriptase (Life Technologies), and incubated at 37°C for 1 h and at 95°C for 5 min. cDNA integrity was assessed by PCR amplification of human β-actin (NC_000007.14). PCR conditions were: 94 °C 1 min, 54° C 1 min, and 72 °C 1 min, for 35 cycles. Human FN-1 (Hs01549976_m1), COL1A1 (Hs01076772_gH), α-SMA (Hs00426835_g1), TGF-β1 (Hs00998133_m1), TIMP-1 (Hs00171558_m1), CAV1 (Hs00971716_m1) and ACTB (Hs99999903_m1) transcripts, were quantified by quantitative real time PCR using TaqMan Universal PCR Master Mix and Assays-on-Demand Gene Expression probes (Applied Biosystems). Reactions were performed in 96-well plates with 25 μl volumes. Cycling conditions were: 2 min at 50 °C, 95 °C for 10 min, and followed by 40 cycles of PCR (15 s at 95 °C and 1 min at 60 °C). Products were detected with the ABI Prism7000 sequence detection system (Applied Biosystems).

The expression of each transcript was normalized to the expression of β-actin and determined as the ratio of the target transcript to the β-actin transcript calculated by 2-ΔΔCt, where ΔΔCt=CtTarget-Ct β-actin. Quantitative PCR data were expressed as “fold changes” relative to control levels.

### Quantitation of soluble/extracellular collagen

Total soluble (non cross-linked) collagen was measured in culture supernatants from control fibroblasts untreated and treated with DMD exosomes, and in culture supernatant from control fibroblasts transfected with miR-199a mimic and inhibitor, by the quantitative dye-binding Sircol assay, according to manufacturer’s instructions.

### TGF-β1 ELISA

TGF-β1 was quantified by ELISA assay (Sigma-Aldrich) following manufacturer’s instructions in control and DMD exosomes, and in control cell culture supernatants before and after exposure of the cells to DMD exosomes.

### Control fibroblasts transfection with miR-199a mimic, inhibitor and scramble

5×10^5^ control fibroblasts were seeded into six-well plates and grown to 60% confluence. Human hsa-miR-199a-5p mimic (assay ID: MC10893), hsa-miR-199a-5p inhibitor (MIMAT0000231, assay ID: MH10893), and control (scramble miRNA) (all from Life Technologies) were allowed to form transfection complexes, with Lipofectamine RNAiMax transfection reagent (Life Technologies) in Opti-MEM I Reduced Serum Medium (Life Technologies) at a final concentration of 30 nM for 15 min at room temperature, and then added to cells. After 24 h, the culture medium was changed to growth medium, and, after further 48 h, cells were either collected for extraction of mRNA and miRNAs, or analysed by immunocytochemistry, and culture media were collected for further analyses. The let-7c/HMGA2 system (High Mobility Group AT-hook 2, Applied Biosystems) was used as positive control of miRNA inhibition experiments, while the miR-1/TWF1 system (Applied Biosystems) was used as positive control for miRNA mimic experiments.

### Exosome loading with miR-199a-5p mimic, inhibitor and scramble

Purified exosomes derived from control fibroblasts were transfected with miR-199a-5p mimic, inhibitor or scramble using Exo-Fect Exosome Transfection Kit (System Biosciences SBI) according with manufacture’s instructions. Briefly, 50 μl of purified exosomes PBS suspension were mixed with 10 μl of Exo-Fect solution and 20 μl of 20 pmol miR-199a mimic, inhibitor or scramble. Transfected exosomes were added to control fibroblasts (1×10^5^ cells/6-wells) and incubated for 24 h. Cell pellets were collected for protein and RNA extraction.

### Animals and experiments on mdx mice

Animal studies were approved by the C. Besta review board, in accordance with Italian Health Ministry guidelines. Animal care and use was in accordance with Italian law and EU directive 2010/63/EU.

The animals were housed in our facility under 12h light/12h dark conditions with free access to food and water. Eight C57BL/6 mice (Jackson Laboratories), 8 weeks old, were injected with 50 μl of 10 μM cardiotoxin (Sigma-Aldrich) in PBS into both tibialis anterior muscles to induce fiber necrosis. DMD and control exosomes (50 μl, corresponding to 70 μg total protein, in PBS) were injected, respectively, into the right lateral tibialis anterior muscle and into the contralateral muscle at 1, 4 and 8 days after cardiotoxin injury. The exosomes injected at 8 days were labelled with the PKH67 Green Fluorescent tag. The total amount of exosomes injected in mice was determined by measuring protein concentration (by DC protein assay) and the protein expression levels of exosome markers Alix and CD63 (by western blot).

Five days after the last injection, the animals were sacrificed by cervical dislocation under anaesthesia; the tibialis anterior muscles were rapidly removed, frozen in isopentane pre-cooled in liquid nitrogen, and maintained in liquid nitrogen pending use. Hematoxylin and eosin (H&E), Gomori modified trichrome, and immunohistochemical staining for collagen I and collagen VI were performed on consecutive 6-8 μm-thick cryosections as previously described in Gibertini et al. (54). Sections were examined under a Zeiss Axioplan2 fluorescence microscope.

### Quantitation of muscle fibrosis

The extent of total connective tissue was determined on sections stained for collagen I and collagen VI, at 20x magnification, using the NIH ImageJ software version 1.51i (http://rsb.info.nih.gov/nih-image/), as described (Gibertini *et al*, 2014). Briefly, from each section, 8 fields were selected for content of fluorescent exosomes (4 fields with more than 10 positive spots and 4 fields with less than 10 positive spots), photographed and digitalized. Using the software, the digitalized images were inverted, and a threshold was applied to the photographs to obtain black and white images with areas positive for collagens in black and negative areas in white. Manual corrections were sometimes applied to eliminate non-muscle/non-fibrosis areas or to add areas not recognized by the software. The area positive for collagens was calculated as a percentage of the entire image, and the mean percentage for each group of animals calculated.

### Statistical analysis

Results were expressed as means and standard deviations (±). Differences between DMD and control cell lines were assessed by the two-tailed Student t-test or one-way ANOVA with post-hoc Dunnett’s test, where appropriate. P values ≤ 0.05 were considered significant.

ELISA data for TGF-β1 were processed using a four-parameter logistic model (Elisaanalysis free software; http://www.elisaanalysis.com/).

The linear regression with interacted dummy variables was employed to assess differences among groups. Coefficients whose associated p-values were less than 0.05 were considered significant.

## Acknowledgments

We thank Dr. Antonello Maruotti of the Dipartimento di Giurisprudenza, Economia, Politica e Lingue Moderne, LUMSA Università, Roma and Centre for Innovation and Leadership in Health Sciences, University of Southampton, Southampton, UK, for help with statistics.

We thank the EuroBioBank and Telethon Network of Genetic Biobanks (GTB12001F to MM) for providing biological samples.

This work was supported by grants for current research from the Italian Ministry of Health, years 2015-2017.

## Author Contributions

SZ and MM conceived and designed the experiments, SZ performed most of the in vitro experiments, SG conceived and performed the in vivo experiments and analysed the data. FB performed electron microscopy studies, CB, AR, SS, CI, PB, LM analysed and interpreted the data. RM and MM handled the funding and supervision. SZ and MM wrote the manuscript. All authors read and approved the final manuscript.

